# Blind spots in global soil biodiversity and ecosystem function research

**DOI:** 10.1101/774356

**Authors:** Carlos A. Guerra, Anna Heintz-Buschart, Johannes Sikorski, Antonis Chatzinotas, Nathaly Guerrero-Ramírez, Simone Cesarz, Léa Beaumelle, Matthias C. Rillig, Fernando T. Maestre, Manuel Delgado-Baquerizo, François Buscot, Jörg Overmann, Guillaume Patoine, Helen R. P. Phillips, Marten Winter, Tesfaye Wubet, Kirsten Küsel, Richard D. Bardgett, Erin K. Cameron, Don Cowan, Tine Grebenc, César Marín, Alberto Orgiazzi, Brajesh K. Singh, Diana H. Wall, Nico Eisenhauer

**Affiliations:** German Centre for Integrative Biodiversity Research (iDiv) Halle-Jena-Leipzig, Leipzig, Germany; Institute of Biology, Martin Luther University Halle Wittenberg, Am Kirchtor 1, 06108 Halle(Saale), Germany; Helmholtz Centre for Environmental Research - UFZ, Department of Soil Ecology; Leibniz-Institut DSMZ-Deutsche Sammlung von Mikroorganismen und Zellkulturen, Braunschweig, Germany; Helmholtz Centre for Environmental Research - UFZ, Department of Environmental Microbiology; Institute of Biology, Leipzig University, Germany; Freie Universität Berlin, Institut für Biologie, Altensteinstr. 6, 14195 Berlin, Germany; Berlin-Brandenburg Institute of Advanced Biodiversity Research (BBIB), Altensteinstr. 34, 14195 Berlin, Germany; Departamento de Biología y Geología, Física y Química Inorgánica, Escuela Superior de Ciencias Experimentales y Tecnología, Universidad Rey Juan Carlos, Calle Tulipán Sin Número, Móstoles 28933, Spain; Departamento de Ecología and Instituto Multidisciplinar para el Estudio del Medio “Ramón Margalef”, Universidad de Alicante, Carretera de San Vicente del Raspeig s/n, 03690 San Vicente del Raspeig, Alicante, Spain; Microbiology, Braunschweig University of Technology, Braunschweig, Germany; Helmholtz Centre for Environmental Research - UFZ, Department of Community Ecology; Institute of Biodiversity, Friedrich Schiller University Jena, Dornburger-Straße 159, 07743 Jena, Germany; School of Earth and Environmental Sciences, The University of Manchester, Manchester, M13 9PT, UK; Department of Environmental Science, Saint Mary’s University, Halifax, NS, Canada; Centre for Microbial Ecology and Genomics, Department of Biochemistry, Genetics and Microbiology, University of Pretoria, Pretoria, South Africa; Slovenian Forestry Institute, Večna pot 2, SI-1000 Ljubljana, Slovenia; Instituto de Ciencias Agronómicas y Veterinarias, Universidad de O’Higgins, Rancagua, Chile; Instituto de Ciencias Ambientales y Evolutivas, Universidad Austral de Chile, Valdivia, Chile; European Commission, Joint Research Centre (JRC), Ispra, Italy; Hawkesbury Institute for the environment, Western Sydney University, Penrith, NSW, 2751, Australia; Global Centre for Land-Based Innovation, Western Sydney University, Penrith, NSW, 2751, Australia; School of Global Environmental Sustainability and Department of Biology, Colorado State University, Fort Collins, CO 80523-1036 USA

**Keywords:** Soil ecology, Macroecology, Soil fauna, Soil microbes, Ecosystem functions

## Abstract

Soils harbor a substantial fraction of the world’s biodiversity, contributing to many crucial ecosystem functions. It is thus essential to identify general macroecological patterns related to the distribution and functioning of soil organisms to support their conservation and governance. Here we identify and characterize the existing gaps in soil biodiversity and ecosystem function data across soil macroecological studies and >11,000 sampling sites. These include significant spatial, environmental, taxonomic, and functional gaps, and an almost complete absence of temporally explicit data. We also identify the limitations of soil macroecological studies to explore general patterns in soil biodiversity-ecosystem functioning relationships, with only 0.6% of all sampling sites having a non-systematic coverage of both biodiversity and function datasets. Based on this information, we provide clear priorities to support and expand soil macroecological research.

## 1. Introduction

Soils harbor a large portion of global biodiversity, including microbes (e.g., Bacteria), micro- (e.g., Nematoda), meso- (e.g., Collembola), and macrofauna (e.g., Oligochaeta), which play critical roles in regulating multiple ecosystem functions and services, including climate regulation, nutrient cycling, and water purification^1–6^. Accordingly, recent experimental^7,8^ and observational^9,10^ studies, based either on particular biomes (e.g., drylands) or local sites, have shown that soil biodiversity is of high importance for the maintenance of multifunctionality (i.e. the ability of ecosystems to simultaneously provide multiple ecosystem functions and services^11^) in terrestrial ecosystems.

Nevertheless, with few exceptions^9,12^, global soil biodiversity-ecosystem function relationships have not yet been studied in depth, with macroecological studies evaluating the patterns and causal mechanisms linking soil biodiversity to soil ecosystem functions only emerging in the last decade^10,13–15^. By comparison, albeit with important limitations^16^, there is a plethora of studies describing the global distribution and temporal patterns of aboveground biodiversity^17^, ecosystems^18^, and biodiversity-ecosystem function relationships^12,19–23^, something that is currently mostly absent (but see^24^) in soil macroecological studies due to the lack of temporally explicit data for soil biodiversity and soil related functions.

Despite the mounting number of soil ecology studies, significant gaps and/or geographic and taxonomic biases exist in our understanding of soil biodiversity^25^. Although the existing gaps in global soil biodiversity data are consistent with gaps in other aboveground biota^16,26,27^, these are further exacerbated when described across specific ecological gradients (e.g., differences across altitudinal gradients) and taxa (e.g., Collembola, Oligochaeta)^28^. Further, and almost nothing is known about the temporal patterns in soil biodiversity at larger spatial scales and across ecosystem types. Identifying and filling these gaps on soil species distributions and functions is pivotal to identify the ecological preferences of multiple soil taxa, assess their vulnerabilities to global change drivers, and understand the causal links between soil biodiversity, ecosystem functions and their associated services^16,29^. Despite growing scientific and political interest in soil biodiversity research^25^, little to no attention has been given to the governance of soil ecosystems (Fig. S1), which has resulted in a lack of inclusion of soil biodiversity and functions in decision making regarding land management debates, conservation, and environmental policy^30^.

In contrast to groups of organisms from other realms (e.g., aboveground terrestrial^31^) for which the Global Biodiversity Information Facility (GBIF) constitutes already the main global data hub^32,33^, soil organisms are poorly represented, with distribution data on soil species spread across the literature and a number of platforms (e.g., the global Ants database^34^, the Earth Microbiome project^35^). Across all available soil biodiversity data, major issues remain regarding the spatial and temporal representativeness (e.g., absent data in most tropical systems) of data, and coverage of taxonomic groups of soil biota (e.g., most focus on fungi and Bacteria), which limits our capacity to comprehensively assess and understand soil systems at multiple temporal and biogeographic scales. More importantly, both the lack of representativeness and the distribution of gaps in global soil biodiversity and ecosystem function research hampers the prioritization of future monitoring efforts^16^. Such a knowledge deficit in soil biodiversity also prevents stakeholders from taking appropriate management actions to preserve and maintain important ecosystem services^36^, such as food and water security, for which soils are the main provider. Therefore, it is both timely and relevant to identify these blind spots in global soil macroecological knowledge and research. By doing so, we can assess their main causes and line up potential solutions to overcome them.

Here, we identify fundamental gaps in soil macroecological research by analysing the distribution of sampling sites across a large range of soil organisms and ecosystem functions. In a review of current literature, we collected sample locations from most existing studies focused on soil macroecological patterns. The studies were then organized according to different soil taxonomic groups and ecosystem functions studied (nine and five categories, respectively, see Methods for more details). Since the mere accumulation of data will not significantly advance ecological understanding^37,38^, it is important to identify how well the current studies cover the range of existing environmental conditions on Earth, including soil properties, climate, topography, and land cover characteristics^39,40^. Finally, we examined how these macroecological studies have captured the diversity of global environmental conditions to identify critical ecological and geographical “blind spots” of global soil ecosystem research (e.g., specific land use types, soil properties, climate ranges; see Methods for more detail). By identifying the environmental conditions that have to be covered in future research and monitoring to draw an unbiased picture of the current state of global soils as well as to reliably forecast their futures, our synthesis goes a significant step beyond recent calls to close global data gaps^25^. Therefore, our comprehensive spatial analysis will help researchers to design future soil biodiversity and ecosystem function surveys, to support the mobilization of existing data, and to inform funding bodies about the allocation of research priorities in this important scientific field.

## 2. Results and Discussion

### 2.1. Biogeographical biases

From our literature search, we collected details on locations of 11,065 individual sampling sites representing studies on soil biodiversity [N=7,631; 68.9% of the total number of sites] and ecosystem functions [N=3,497; 31.6% of the total number of sites] (Fig. 1). Bacteria, fungi, and soil respiration (Fig. 1a) were the best-represented soil taxa and functions in our literature survey, respectively. The total number of sites across all studies is quite low when compared with many aboveground macroecological databases that can individually surpass the numbers found here (e.g., the PREDICTS database^41^ contains ~29,000 sites across the globe).

**Fig. 1.**
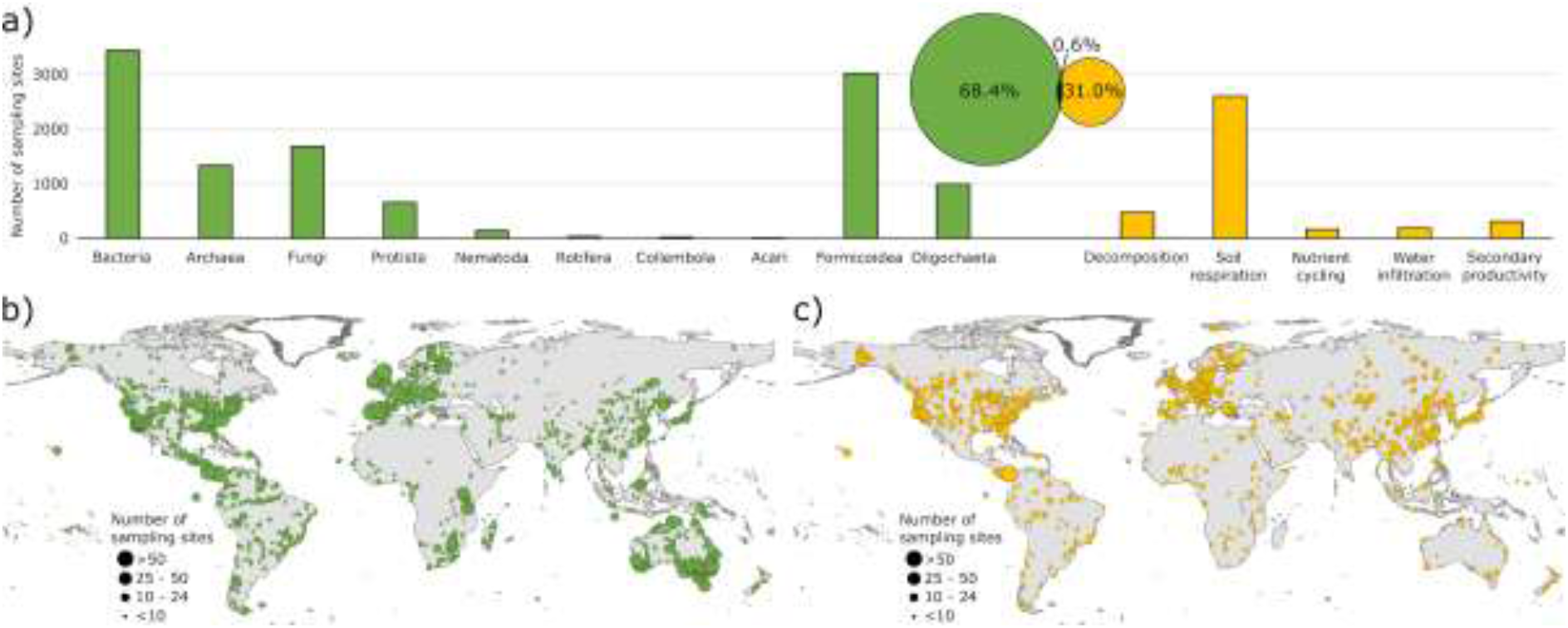
Global distribution of sampling sites for soil taxa and soil ecosystem functions. (a) corresponds to the global number of individual sampling sites for each soil taxon (on the left - in green) and ecosystem function (on the right - in orange). The venn diagram indicates the proportion of sampling sites for soil taxa (in green) and functions (in orange), and the 0.6% [N=63] of overlap between biodiversity and function data points (this number does not mean that soil biodiversity and function were assessed in the same soil sample or during the same sampling campaign; i.e., there could still be a thematic or temporal mismatch), relative to the total number of sampling sites covered by the studies. The maps show the overall spatial distribution of sampling sites for all taxa (b) and soil ecosystem functions (c). The size of the circles corresponds to the number of sampling sites within a 1-degree grid ranging from <10 to >50.

Globally, soil biodiversity and ecosystem function data are not evenly distributed. Bacteria [N=3,453], fungi [N=1,687], and Formicoidea [N=3,024] (which together concentrate 71.6% of all soil biodiversity records) have comparatively large and geographically balanced distributions when compared to Nematoda [N=149], Rotifera [N=41], Collembola [N=27], and Acari [N=10], which have a substantially lower number of sampling sites and more scattered distributions (see Fig. S2a for more detail). In the case of Bacteria and fungi, the relatively high number of sampling sites reflects a community effort to assemble databases based on collections from different projects^10,42^. In the case of Formicoidea, the availability of data reflects the outcome of systematic global sampling initiatives^43^ or a combination of both^44^.

Soil ecosystems are by nature very heterogeneous at local scales^45^. Having a small and scattered number of sampling sites, for both soil functions and taxa, limits the power of current global analyses to evaluate macroecological relationships between soil biodiversity and ecosystem function, particularly for nutrient cycling and secondary productivity, which have strong local inter-dependencies^46^. In fact, from the five functions assessed here, there is a clear concentration of studies on soil respiration, accounting for 69.1% [N=2,616] of all function records (see Fig. S2b for more detail). Thus, our study provides evidence for a lack of matching data for soil biodiversity and multiple ecosystem functions in current global datasets. Due to the dependency of these and other soil functions on biodiversity^2,47^, being able to deepen our understanding of the strength and distribution of expected biodiversity and ecosystem function relationships is critical to better inform management and policy decisions^48^. In this context, only 0.6% of all sampling sites have an overlap between biodiversity and function datasets (corresponding to 63 sampling sites), with a non-systematic coverage of just a few taxa and functions across sites. Nowadays, macroecological studies on aboveground biodiversity and ecosystem functioning^19,41,49–52^ rely on data mobilization mechanisms that allow for data to be often reused to address multiple research questions. By contrast, apart from some taxonomic groups (i.e., Bacteria and fungi) soil macroecological studies based on observational data have a very small degree of overlap and still remain conditioned by poor data sharing and mobilization mechanisms^53–55^.

We also discovered that most studies are based on single sampling events, i.e., without repeated measurements in time for the same sampling sites. Being able to study how communities and functions change over time is essential for assessing trends in key taxa and functions, and their vulnerability to global change^17^. Our global survey suggests that such information is almost nonexistent in large-scale soil biodiversity and ecosystem functions studies. Thus, for most soil communities and functions, although local studies exist^56,57^, understanding the global trends and the implications of global change drivers and scenarios is difficult and limited by the absence of globally distributed and temporally explicit observational data.

### 2.2. Ecological blind spots

Overall, both soil biodiversity and ecosystem function variables reveal a high degree of spatial clustering across global biomes: temperate biomes (especially broadleaved mixed forests and Mediterranean) contain more sampling sites than tundra, flooded grasslands and savannas, mangroves, and most of the tropical biomes, with the exception of moist broadleaf forests (Fig. S3). This spatial clustering is even more pronounced in studies of ecosystem functions, with temperate systems being overrepresented with 62% of all sampling sites, while the rest of the globe has scattered information on soil conditions. This likely reflects differences in funding availability and research expertise across countries^27,58^. In fact, for taxa like Collembola and Nematoda, most of sampling sites are concentrated in temperate regions, with very few being documented in other regions. Further, the availability of soil biodiversity and function data is especially scarce, and in some cases non-existent: in tropical and subtropical regions (see Fig. S3 for more details), which are among the most megadiverse places on Earth, montane grasslands, and deserts. In many cases, local experts may exist, although their contributions are often not included in macroecological studies. At the same time, for many of the best-represented regions in the globe, there is rarely a complete coverage of soil taxa and functions, with records often being overinflated by one or two densely sampled taxa (e.g., Bacteria and fungi) or functions (e.g., soil respiration).

The range of environmental conditions currently described within soil macroecological studies is critical to understand the relationship between soil biodiversity, ecosystem functions, and key environmental conditions (e.g., the known relationship between Bacteria richness and pH^59^ or the dependence of soil respiration on temperature^60,61^). In this context, the complete range of soil carbon levels existing on Earth is not well covered, with soils of very high and low carbon contents (Fig. 2a) being underrepresented compared with their global distribution. The same applies to soil type, with only a fraction of soil types being well covered (i.e., acrisols, andosols, cambisols, kastanozems, luvisols and podzols), while others are significantly underrepresented or completely absent (e.g., durisols, stagnosols, umbrisols; Fig. 2o). In contrast, our study identified over- and underrepresented environmental conditions in soil biodiversity and function studies (Fig. 2). For example, some soil properties are well represented across studies, such as soil texture (i.e., sand, silt, and clay content) and pH, with the exception of extreme ranges (e.g., pH > 7.33 or silt content < 19%).

**Fig. 2.**
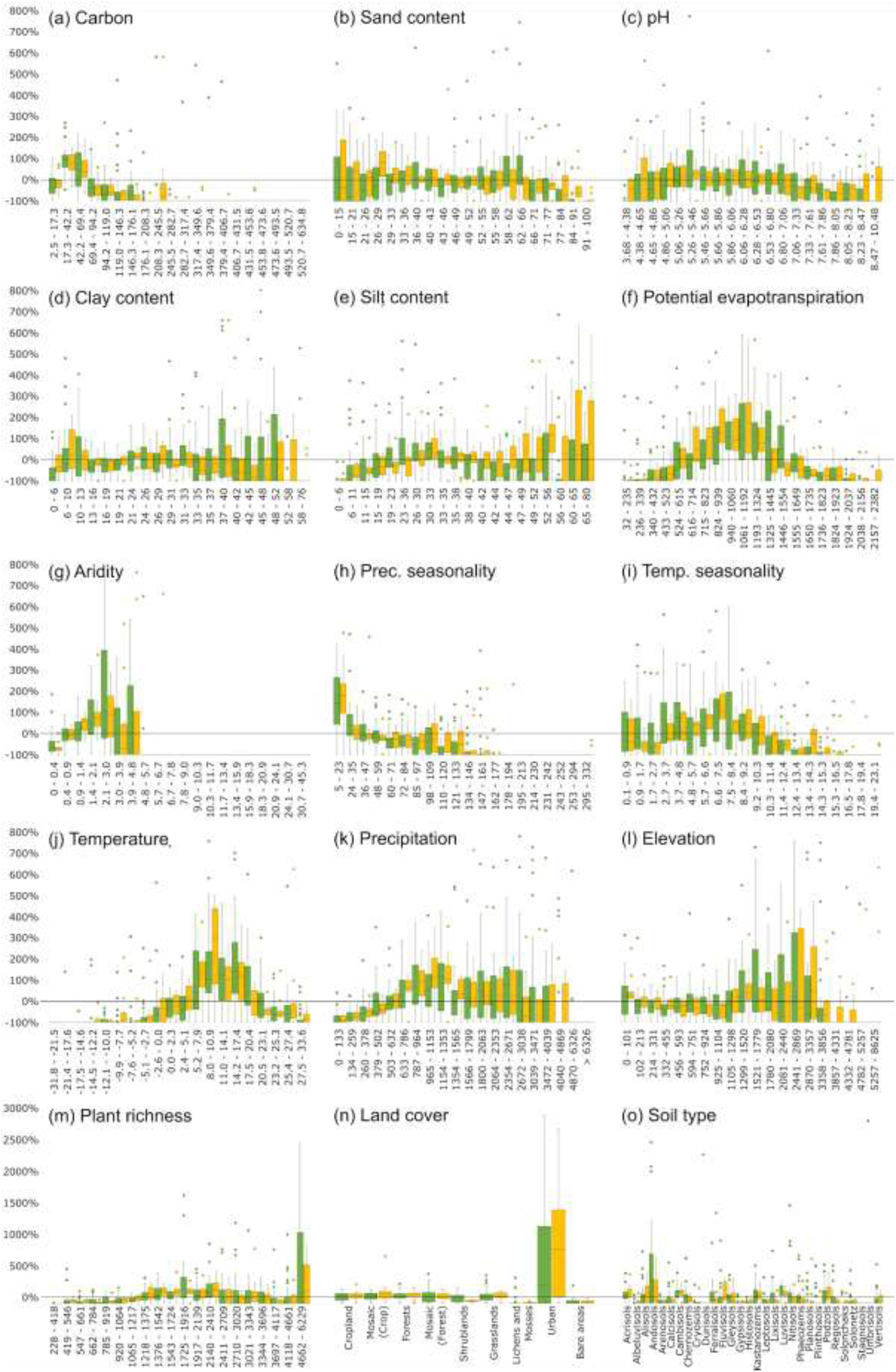
Global soil ecological blind spots. Values (y-axis) correspond to the percentage of sites per study per class when compared with the global proportional distribution of each class within each variable defining soil ecosystems (soil biodiversity studies in green and ecosystem function studies in orange): (a) soil carbon [g/soil kg]^65^; (b) sand content [%]^65^; (c) soil pH^65^; (d) clay content (%)^65^; (e) silt content (%)^65^; (f) potential evapotranspiration^66^; (g) aridity index^66^; (h) precipitation seasonality^67^; (i) temperature seasonality^67^; (j) mean annual temperature^67^; (k) mean total precipitation^67^; (l) elevation^68^; (m) vascular plant richness^69^; (n) land cover^70^; (o) soil type^65^. The zero black line corresponds to a situation where the proportion of sites in a given class within a study matches the global proportional representation of the same class. Although outliers were not eliminated, for representation purposes these were omitted >800% between panels a to l and <3000% for panels m to o.

In contrast to to soil conditions, climate variability is systematically poorly covered in soil biodiversity and function studies, with significant climatic ranges being almost completely missing (Fig. 2f-k). These include low and high potential evaporation and aridity areas, areas with high climate seasonality, low precipitation and extreme temperatures (i.e., very hot and very cold systems), with no overall significant differences between biodiversity and ecosystem function studies. Drylands, for example, cover ~45% of the land surface^62^ and have been shown to be highly diverse in terms of soil biodiversity and with strong links to specific ecosystem functions^24,63^, but are often underrepresented. Climatic conditions (current and future) have strong influences on both soil organisms^57^ and functions^60,64^; as such, assessing a wide range of these conditions, including climatic extremes, is fundamental to describe the complex dynamics of soil systems. This issue is further exacerbated when looking at specific climate combinations (Fig. 3c), where 59.6% of the global climate is not covered by any of the studies considered.

**Fig. 3.**
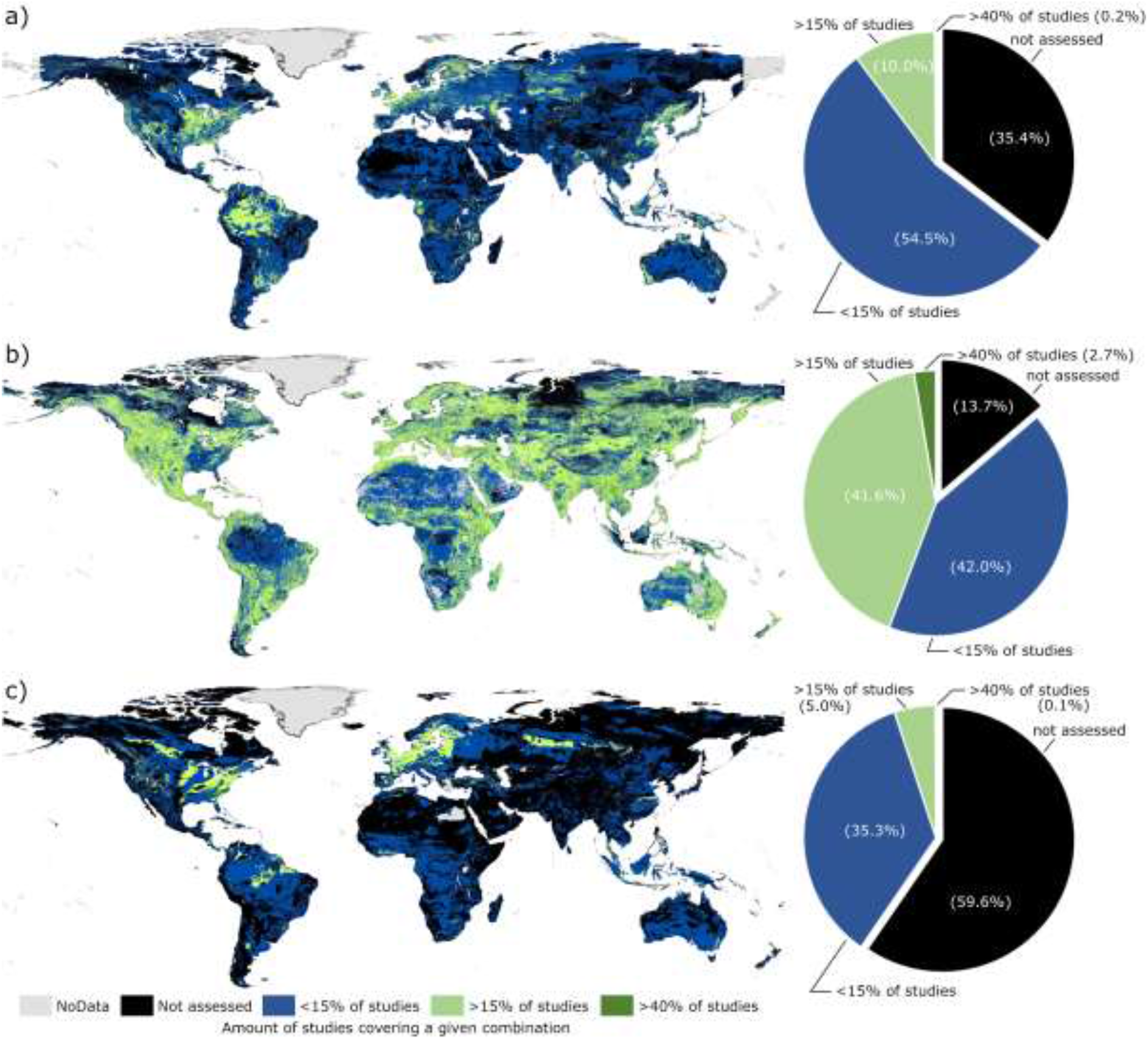
The extent to which main soil environmental characteristics are assessed across macroecological studies. Colours correspond to the amount of studies covering a given combination of characteristics (see Methods for more details) within: a) land cover (including the combination of land cover, plant diversity and elevation); b) soils (including the combination of organic carbon content, sand content and pH); and c) climate (including the combination of mean precipitation and temperature, and their seasonality). Black corresponds to combinations that were not assessed by any of the studies here included; in blue are the combinations assessed by less than 15% of the studies (N= 7); in light green the variable combinations assessed by less than 40% of the studies (N=18); and in dark green, the variable combinations assessed by more than 40% of the studies. All combinations were created by a spatial overlap using the same class distribution of each variable as in Fig. 2 (see Methods and Fig. S4a-e for more details).

Although representing a major driver of soil biodiversity and function^4^, land-cover based studies have shown different responses across groups of soil organisms^56,71,72^ and specific functions^73,74^. While, in general, land cover types are well covered, sites in the proximity of urban areas are disproportionately represented (Fig. 2n). Lichens, mosses, and bare areas have been neglected, and shrublands are not well represented in ecosystem function assessments. These gaps may have important implications, particularly when they correlate with understudied ecosystems like drylands or higher latitude systems that may harbor high biodiversity^63^, but for which patterns are mostly unknown. In this context, the present analysis indicates that low diversity areas (here represented as plant richness^69^) are absent from most studies or poorly represented, with the focus being mostly on higher diversity areas. Concurrently, it has been suggested that there may be important mismatches between above- and belowground biodiversity across the globe^75^, i.e., there are huge areas where aboveground biodiversity does not well predict belowground biodiversity.

When looking at how belowground studies cover the combinations of aboveground diversity (Fig. 3a) and of soil conditions (Fig 3b), important mismatches are observed. We also looked at combinations of environmental gradients. Here, although most soil-related environmental combinations (Fig. 3b) are well covered across studies, the same does not apply when looking at the aboveground diversity (Fig. 3a), which shows a very good coverage in forest and crop areas with above average plant richness in mid to low elevations, while other environmental combinations are underrepresented. Overall, while it is unreasonable to expect all macroecological studies to cover all possible soil conditions, the systematic underrepresentation of many soil characteristics observed here may undermine our capacity to generalise results given that they do not capture the full ecological space of soil organisms.

Many of the reasons and drivers of existing data gaps have already been illustrated in recent literature for aboveground systems^16^ (e.g., accessibility, proximity to large cities, etc.). In the case of soil biodiversity and ecosystem functions, these blind spots are further reinforced because of the lack of standardized protocols for acquiring biodiversity and ecosystem function data. This translates into an absence of comparable data, which is even more pronounced than in other systems^16,76^. Nevertheless, there is a continuous movement towards improving data mobilization and international collaborations that could help overcome these issues if steered in the direction of underestimated taxa and/or functions identified here^77^.

In a changing world where soil biodiversity shifts are being systematically reported^78–80^, and where current forecasts are pointing to increases in land-use intensity^81,82^, desertification^83^, and rapid climate change^84–87^, understanding if and to what extent biodiversity changes are happening in soil communities is of high importance. This is particularly relevant to assess causal effects between changes in biodiversity and ecosystem function (e.g., are changes in biodiversity occurring because of changes in function, paired with them, or despite them, and *vice versa*), which is even more relevant if key ecosystem functions (e.g., carbon sequestration) are the subject of evaluation.

Filling the knowledge gap on large-scale temporal trends in soil biodiversity and ecosystem function cannot be achieved without spatially explicit studies based on resampled locations. This could be done with a proper global monitoring framework that is recognized and supported by a large number of countries, which currently does not exist. In this context, given the strength of recognized soil taxa interactions^88^, biodiversity and ecosystem function relationships^24^, and above-belowground interactions^89^, these large-scale monitoring activities and research studies should consider going beyond traditional single taxa/function approaches and collect information on the multiple dimensions of soil ecosystems^28^, while at the same time expanding/supporting surveys to cover the blind spots of soil macroecological research (Fig. 3).

### 2.3. Challenges to move beyond blind spots

Across all soil taxa and functions, the geographical and ecological blind spots identified here often emerge from a number of obstacles specific to soil ecology^77^ (see summary in Table 1). Soil macroecologists face many challenges and constraints spanning from a lack of methodological standards and scientific expertise in different taxonomic groups^90–92^, to limitations caused by the current implementation of the Convention for Biological Diversity (CBD) and the Nagoya Protocol^93,94^. While the first has more immediate, albeit non-trivial solutions (e.g., by expanding the language pool of the researchers and studies included^16,95^ and by applying common standards for sampling, extraction, and molecular protocols^96–99^), the latter contains systemic issues that go beyond soil ecology alone. In this context, although the CBD and the Nagoya Protocol were created to protect countries while making the transfer of biological material more agile, numerous states have either not yet implemented effective national Access and Benefit Sharing (ABS) laws or have implemented very strict regulations^100,101^. Yet, even after 25 years of the CBD and the ABS framework being in place, the major motivation for a strict national regulation - the anticipated commercial benefits and high royalties from the “green gold” - has not yet materialized^93,102^.

**Table 1.** Summary of the main obstacles soil ecologist face to create a global soil biodiversity monitoring network and the priority actions to overcome them.

| Challenges | Priority Actions |  |  |
| --- | --- | --- | --- |
|  | Researchers | Institutions | Policymakers |
| Legal issues regarding the transport and sharing of soil samples and biological data | ● Raise the awareness of institutions and decision-makers about the importance of these legal bottlenecks for the development of | Develop a legal understanding of the implications of material transfer mechanisms for soil samples and provide support to | Establish global multilateral solutions and International Treaties focused on soil biodiversity and ecosystem |
|  | international research programs. | researchers also by promoting knowledge and expertise exchange.<br>Support and facilitate the establishment of international consortia and bilateral institutional agreements particularly with developing countries | function research.<br>Establish knowledge transfer mechanisms for soil-related research together with the classification of soil samples for research purposes. |
| Scattered literature and lack of mobilization/ systematization of local studies | Invest in data harmonization, synthesis, meta-analysis approaches, data collation, and proper metadata to improve currently available datasets (e.g., through GBIF for soil biodiversity).<br>Define and publish data standards that allow for better data transfer focussing on the methods, reporting in standard units, and best practices for data availability.<br>Increase the focus on understudied soil groups (e.g., collembola, acari, protists, mammals) and functions (e.g., soil aggregate stability, bioturbation, nutrient cycling).<br>Establish effective coordination of current networks to support the development of integrated ecological assessments of the soil realm | Adoption of available data and methods standards<sup>99,103–107</sup> and support the establishment and maintenance of data repositories and open access policies. | Support open access partnerships (e.g., the German DEAL<sup>108</sup>) to facilitate knowledge transfer and collaboration across countries and researchers from different backgrounds and expertise.<br>Improve the digitally available data on soil biodiversity and ecosystem function by supporting the expansion of current global databases (e.g., GBIF) or the creation of interoperable data infrastructures on soil function data. |
| Lack of temporally explicit information on soil biodiversity and functions | Identify relevant sites - e.g., sites covering a wide range of taxa or functions and/or a high degree of standardization - for resampling.<br>Revisit already sampled sites to obtain temporal measurements of soil biodiversity and ecosystem function. | Institutional support of long-term databases and collections of soils, soil functional data, and soil biological material. | Create funding schemes for strategic long-term research projects on soil monitoring and research (e.g., using the LTER framework as an example<sup>109</sup>). |
| Lack of globally distributed expertise, research funding and infrastructures | Promote knowledge transfer mechanisms and capacity building, especially with developed countries that might see little advantage of being involved in a global network that only offer co-authorship as the main benefit.<br>Setup international workshops, summer schools, or classes with a focus on educating the next generation of scientists on different aspect of soil ecology. | Build on or expand current networks to include knowledge transfer activities, namely on education, methods calibration, sharing research facilities, and taxonomic expertise. | Promote funding flexibility to train and empower researchers across countries and/or regions, also allowing local scientists, particularly in the developing world, to conduct soil biodiversity and ecosystem function research.<br>Establish soil health as a research priority beyond farming areas and with a special focus on ecological conservation of soil organisms and ecosystem functions. |

Researchers have yet to coordinate a global effort to characterize the multiple aspects of soil biodiversity and function in a comprehensive manner, with the current literature being dominated by scattered, mostly local studies focused on specific soil organisms and/or functions. Although here we do not assess the potential of local studies to overcome the current blind spots, other studies^34,35,64^ have shown that, with a significant effort in standardization and data mobilization, local and regional studies add fundamental knowledge and empower local researchers to participate in global initiatives. In fact, several studies not included in this assessment can provide a finer-scale resolution in many areas of the globe^71,110,111^.

Nevertheless, their spatial extent systematically coincides with overrepresented areas (e.g., temperate areas), and their taxonomic and functional focus is mostly on the already prevailing taxa (i.e., Bacteria and fungi) and functions (i.e., soil respiration), potentially increasing existing biases. This increases the relevance of facilitating data mobilization from regions and, more importantly, environmental conditions that are systematically not covered by macroecological studies.

In parallel, and given the nature of global change drivers, understanding their influence on local soil communities and ecosystem functioning requires global macroecological approaches that can provide context, predictions, and concrete suggestions to policymakers across the globe. Yet these macroecological approaches will be less effective in providing relevant outputs at national scales if they are based on data extrapolated from other countries; they would be strongly improved if local data were made available^25,112^. Without more comprehensive studies seeking answers to large-scale soil ecological questions - often involving dealing with multiple scales (temporal and spatial) and a number of thematic and taxonomic depths^75^ - it is difficult to deepen soil macroecological knowledge^113^. This is particularly relevant in testing biodiversity and ecosystem function relationships at the global scale, or trying to address specific societal issues (e.g., the attribution of climate and land-use change as drivers of soil ecological change or general biodiversity trends)^17^.

Another major challenge is associated with the fact that currently ABS agreements are bilateral. This hinders global soil ecology initiatives as it requires that providers and receivers need limited individual material transfer agreements. Thus, for a global initiative, this can amount to hundreds of material transfer agreements. However, there is an increasing quest for global solutions and multilateral systems, such as the International Treaty on Plant Genetic Resources for Food and Agriculture (IT PGRFA; www.fao.org/3/a-i0510e.pdf). or other harmonized best practices examples like the Global Genome Biodiversity Network^94^. As the commercial value of soil organisms is regarded to be zero *in sit*u^114^, and as these are mostly ubiquitously distributed at the highest taxonomic level (e.g., Bacteria and fungi), soil *per se* has no commercial value as it does not match the criteria that “provider countries host unique and unmatched biodiversity” of the Nagoya Protocol^115^. Therefore, a global multilateral solution, similar to the examples listed above (e.g., the IT PGRFA), but focused on facilitating the exchange of soil samples to drive basic research on soil biodiversity, taxonomy, and ecosystem functioning, while still safeguarding against the spread of foreign genotypes, is pressingly needed. At the same time, as long as bilateral ABS agreements are required, researchers should engage with local policymakers to enable unrestricted soil biodiversity and ecosystem function research, as was the case for Brazil from 2006 to 2016^116^.

### 2.4. Looking for solutions to unearth global observations

Globally, soil habitats are under constant pressure from major threats, such as climate change, land use change and intensification, desertification, and increased levels of pollution. Here, we argue for a global monitoring initiative that systematically samples soil biodiversity and ecosystem functions across space and time. Such a global initiative is urgently needed to fully understand the consequences of ongoing global environmental change on the multiple ecosystem processes and services supported by soil organisms (Table 1). This requires that current and future funding mechanisms include higher flexibility for the involvement of local partners from different countries in global research projects. Given that soil ecological research requires cross-border initiatives^77^ and expensive infrastructure, there is a need for flexible funding with proper knowledge transfer mechanisms to sustain global soil macroecological research. Such knowledge will in turn contribute to advancing our understanding of macroecological patterns of soil biodiversity and ecosystem function, thereby fulfilling national and global conservation goals^114,117,118^.

Considering the current pool of literature, improving the digitally available data on soil biodiversity and ecosystem function should be a top priority that could be made possible by systematically mobilizing the underlying data^119^ in already existing open access platforms (e.g., GBIF). Achieving this goal on shared knowledge and open access data will return benefits beyond making global soil biodiversity surveys possible. It will allow local researchers to expand their own initiatives, create a more connected global community of soil ecologists, bypassing publication and language limitations, and potentially open doors in countries that may otherwise be reluctant in sharing their soil biodiversity data^27^.

In parallel, coordinated sampling strategies based on standardized data collection and analysis are needed to improve soil macroecological assessments. From our results, it is clear that most, if not all, studies look at only a fraction of the soil realm without much spatial and thematic complementarity of global environmental conditions. Also, the significantly small overlap between biodiversity and functional studies indicates that most community assessments disregard the ecosystem functions that these provide and *vice versa*, prompting a call for more complex approaches that can show potential links and global ecosystem services. Our study helps to identify global target locations and biomes which need to be given priority in future surveys. Future sampling strategies would greatly benefit from coordinated sampling campaigns with biodiversity and function assessments at the same locations and ideally from the same soil samples to improve the current spatial-temporal resolution of data on soil biodiversity and ecosystem functions.

These two complementary pathways (i.e., data mobilization and sharing of current literature and a globally standardized sampling) if done in a spatially explicit context, and following standardized protocols, could ultimately inform predictive modelling frameworks for soil ecosystems to track the fulfilment of global/national biodiversity targets, policy support, and decision making. Taken together, our study shows important spatial and environmental gaps across different taxa and functions that future macroecological research should target, and a need to collect temporal datasets to explore if current aboveground biodiversity declines are also seen in belowground taxa. With the identification of global spatial, taxonomic, and functional blind spots, and the definition of priority actions for global soil macroecological research^75^, our synthesis highlights the need for action to facilitate a global soil monitoring system that overcomes the current limitations.

## Acknowledgements

This manuscript developed from discussions within the German Centre of Integrative Biodiversity Research funded by the German Research Foundation (DFG FZT118). CAG and NE acknowledge funding by iDiv (DFG FZT118) Flexpool proposal 34600850. CAG, AHB, JS, AC, NGR, SC, LB, MCR, FB, JO, GP, HRPP, MW, TW, KK and NE acknowledge funding by iDiv (DFG FZT118) Flexpool proposal 34600844. N.E. acknowledges funding by the DFG (FOR 1451) and the European Research Council (ERC) under the European Union’s Horizon 2020 research and innovation programme (grant agreement no. 677232).

## Authors contributions

CAG, AHB, JS, AC, NGR, SC, LB, MCR, FB, JO, GP, HRPP, MW, TW, KK, FTM and NE conceived and designed the study; CAG, AHB, JS, AC, NGR, SC, LB, MCR, FB, JO, GP, HRPP, MW, TW, KK, FTM, EC, CM, AO, DHW, and NE contributed to the data acquisition and all authors had substantial contributions to the interpretation of the data, writing and review of the final manuscript.

## References

1. Wall, D. H. et al. Soil ecology and ecosystem services. 406 (Oxford University Press, 2012).

2. Blouin, M. et al. A review of earthworm impact on soil function and ecosystem services. Eur. J. Soil. Sci. 64, 161–182 (2013).

3. Baveye, P. C., Baveye, J. & Gowdy, J. Soil ‘Ecosystem’ Services and Natural Capital: Critical Appraisal of Research on Uncertain Ground. Front. Environ. Sci. Eng. China 4, 1–49 (2016).

4. Bardgett, R. D. & van der Putten, W. H. Belowground biodiversity and ecosystem functioning. Nature 515, 505–511 (2014).

5. Heemsbergen & Hal, V. Biodiversity Effects on Soil Processes Explained by Interspecific Functional Dissimilarity Biodiversity Effects on Soil Processes Explained by Interspecific. Science 306, 8–10 (2004).

6. Reich, P. B. et al. Impacts of biodiversity loss escalate through time as redundancy fades. Science 336, 589–592 (2012).

7. Schuldt, A. et al. Biodiversity across trophic levels drives multifunctionality in highly diverse forests. Nat. Commun. 9, 2989 (2018).

8. Risch, A. C. et al. Size-dependent loss of aboveground animals differentially affects grassland ecosystem coupling and functions. Nat. Commun. 9, 3684 (2018).

9. Soliveres, S. et al. Biodiversity at multiple trophic levels is needed for ecosystem multifunctionality. Nature 536, 456–459 (2016).

10. Maestre, F. T. et al. Increasing aridity reduces soil microbial diversity and abundance in global drylands. Proceedings of the National Academy of Sciences 112, 201516684 (2015).

11. Manning, P. et al. Redefining ecosystem multifunctionality. Nature Ecology & Evolution 2, 427–436 (2018).

12. Maestre, F. T. et al. Plant species richness and ecosystem multifunctionality in global drylands. Science 335, 214–218 (2012).

13. Song, D. et al. Large-scale patterns of distribution and diversity of terrestrial nematodes. Appl. Soil Ecol. 114, 161–169 (2017).

14. Delgado-Baquerizo, M. et al. Palaeoclimate explains a unique proportion of the global variation in soil bacterial communities. Nat Ecol Evol 1, 1339–1347 (2017).

15. Pärtel, M., Bennett, J. A. & Zobel, M. Macroecology of biodiversity: disentangling local and regional effects. New Phytol. 211, 404–410 (2016).

16. Meyer, C., Kreft, H., Guralnick, R. & Jetz, W. Global priorities for an effective information basis of biodiversity distributions. Nat. Commun. 6, 8221 (2015).

17. Dornelas, M. et al. Assemblage time series reveal biodiversity change but not systematic loss. Science 344, 296–299 (2014).

18. Hansen, M. C. et al. High-resolution global maps of 21st-century forest cover change. Science 342, 850–853 (2013).

19. Grace, J. B. et al. Integrative modelling reveals mechanisms linking productivity and plant species richness. Nature 529, 390–393 (2016).

20. Liang, J. et al. Positive biodiversity-productivity relationship predominant in global forests. Science 354, (2016).

21. Duffy, J. E., Godwin, C. M. & Cardinale, B. J. Biodiversity effects in the wild are common and as strong as key drivers of productivity. Nature 549, 261–264 (2017).

22. van der Plas, F. et al. Jack-of-all-trades effects drive biodiversity-ecosystem multifunctionality relationships in European forests. Nat. Commun. 7, 11109 (2016).

23. van der Plas, F. et al. Continental mapping of forest ecosystem functions reveals a high but unrealised potential for forest multifunctionality. Ecol. Lett. 21, 31–42 (2018).

24. Delgado-Baquerizo, M. et al. Microbial diversity drives multifunctionality in terrestrial ecosystems. Nat. Commun. 7, 10541 (2016).

25. Cameron, E. K. et al. Global gaps in soil biodiversity data. Nat Ecol Evol l2, 1042–1043 (2018).

26. Wetzel, F. T. et al. Unlocking biodiversity data: Prioritization and filling the gaps in biodiversity observation data in Europe. Biol. Conserv. 221, 78–85 (2018).

27. Amano, T. & Sutherland, W. J. Four barriers to the global understanding of biodiversity conservation: wealth, language, geographical location and security. Proc. Biol. Sci. 280, 20122649 (2013).

28. Eisenhauer, N., Bonn, A. & Guerra, C. A. Recognizing the quiet extinction of invertebrates. Nat. Commun. (2019).

29. Pereira, H. M., Navarro, L. M. & Martins, I. S. Global Biodiversity Change: The Bad, the Good, and the Unknown. Annu. Rev. Environ. Resour. 37, 25–50 (2012).

30. Paleari, S. Is the European Union protecting soil? A critical analysis of Community environmental policy and law. Land use policy 64, 163–173 (2017).

31. Meyer, C., Weigelt, P. & Kreft, H. Multidimensional biases, gaps and uncertainties in global plant occurrence information. Ecol. Lett. 19, 992–1006 (2016).

32. Costello, M. J., Michener, W. K., Gahegan, M., Zhang, Z.-Q. & Bourne, P. E. Biodiversity data should be published, cited, and peer reviewed. Trends Ecol. Evol. 28, 454–461 (2013).

33. Bingham, H. C., Doudin, M. & Weatherdon, L. V. The biodiversity informatics landscape: elements, connections and opportunities. (2017).

34. Gibb, H. et al. A global database of ant species abundances. Ecology 98, 883–884 (2017).

35. Thompson, L. R. et al. A communal catalogue reveals Earth’s multiscale microbial diversity. Nature 551, 457–463 (2017).

36. de Groot, R. S., Alkemade, R., Braat, L., Hein, L. & Willemen, L. Challenges in integrating the concept of ecosystem services and values in landscape planning, management and decision making. Ecol. Complex. 7, 260–272 (2010).

37. Stewart, G. Meta-analysis in applied ecology. Biol. Lett. 6, 78–81 (2010).

38. Rillig, M. C. et al. Biodiversity research: data without theory—theory without data. Frontiers in Ecology and Evolution 3, 20 (2015).

39. Coleman, D. C., Callaham, M. A. & Crossley, D. A., Jr. Fundamentals of Soil Ecology. (Academic Press, 2017).

40. Lavelle, P. & Spain, A. Soil Ecology. (Springer Science & Business Media, 2001).

41. Hudson, L. N. et al. The database of the PREDICTS (Projecting Responses of Ecological Diversity In Changing Terrestrial Systems) project. Ecol. Evol. 7, 145–188 (2017).

42. Tedersoo, L. et al. Global diversity and geography of soil fungi. Science 346, 1256688 (2014).

43. Orgiazzi, A. et al. Global soil biodiversity atlas. (2016).

44. Guenard, B., Weiser, M. D. & Gomez, K. The Global Ant Biodiversity Informatics (GABI) database: synthesizing data on the geographic distribution of ant species (Hymenoptera: Formicidae). Myrmecol. News (2017).

45. Nielsen, U. N. et al. The enigma of soil animal species diversity revisited: the role of small-scale heterogeneity. PLoS One 5, e11567 (2010).

46. Lavelle, P. et al. Soil invertebrates and ecosystem services. Eur. J. Soil Biol. 42, (2006).

47. Evans, T. A., Dawes, T. Z., Ward, P. R. & Lo, N. Ants and termites increase crop yield in a dry climate. Nat. Commun. 2, 262–267 (2011).

48. Eisenhauer, N., Bowker, M. A., Grace, J. B. & Powell, J. R. From patterns to causal understanding: Structural equation modeling (SEM) in soil ecology. Pedobiologia 58, 65–72 (2015).

49. Isbell, F. et al. Biodiversity increases the resistance of ecosystem productivity to climate extremes. Nature 526, 574–577 (2015).

50. Craven, D. et al. Multiple facets of biodiversity drive the diversity-stability relationship. Nat Ecol Evol 2, 1579–1587 (2018).

51. Fraser, L. H. et al. Plant ecology. Worldwide evidence of a unimodal relationship between productivity and plant species richness. Science 349, 302–305 (2015).

52. Newbold, T. et al. Global effects of land use on local terrestrial biodiversity. Nature 520, 45–50 (2015).

53. Borgman, C. L., Wallis, J. C. & Enyedy, N. Little science confronts the data deluge: habitat ecology, embedded sensor networks, and digital libraries. Int J Digit Libr 7, 17–30 (2007).

54. Hampton, S. E. et al. Big data and the future of ecology. Front. Ecol. Environ. 11, 156–162 (2013).

55. Pey, B. et al. Current use of and future needs for soil invertebrate functional traits in community ecology. Basic Appl. Ecol. 15, 194–206 (2014).

56. Tsiafouli, M. A. et al. Intensive agriculture reduces soil biodiversity across Europe. Glob. Chang. Biol. 21, 973–985 (2015).

57. Blankinship, J. C., Niklaus, P. A. & Hungate, B. A. A meta-analysis of responses of soil biota to global change. Oecologia 165, 553–565 (2011).

58. Wheeler, Q. D., Raven, P. H. & Wilson, E. O. Taxonomy: impediment or expedient? Science 303, 285 (2004).

59. Delgado-Baquerizo, M. & Eldridge, D. J. Cross-Biome Drivers of Soil Bacterial Alpha Diversity on a Worldwide Scale. Ecosystems 1–12 (2019).

60. Hursh, A. et al. The sensitivity of soil respiration to soil temperature, moisture, and carbon supply at the global scale. Glob. Chang. Biol. 23, 2090–2103 (2017).

61. Wang, Q., Liu, S. & Tian, P. Carbon quality and soil microbial property control the latitudinal pattern in temperature sensitivity of soil microbial respiration across Chinese forest ecosystems. Glob. Chang. Biol. 24, 2841–2849 (2018).

62. Prăvălie, R. Drylands extent and environmental issues. A global approach. Earth-Sci. Rev. 161, 259–278 (2016).

63. Delgado-baquerizo, M. et al. A global atlas of the dominant bacteria found in soil. Science 325, 320–325 (2018).

64. Djukic, I. et al. Early stage litter decomposition across biomes. Sci. Total Environ. (2018).

65. Hengl, T. et al. SoilGrids1km--global soil information based on automated mapping. PLoS One 9, e105992 (2014).

66. Trabucco, A., Zomer, R. J., Bossio, D. A., van Straaten, O. & Verchot, L. V. Climate change mitigation through afforestation/reforestation: A global analysis of hydrologic impacts with four case studies. Agric. Ecosyst. Environ. 126, 81–97 (2008).

67. Karger, D. N. et al. Climatologies at high resolution for the Earth land surface areas. Scientific data 1–19 (2017).

68. Global Multi-resolution Terrain Elevation Data 2010 (GMTED2010) | The Long Term Archive. Available at: https://lta.cr.usgs.gov/GMTED2010. (Accessed: 6th December 2018)

69. Kreft, H. & Jetz, W. Global patterns and determinants of vascular plant diversity. Proc. Natl. Acad. Sci. U. S. A. 104, 5925–5930 (2007).

70. European Space Agency. ESA - Land Cover CCI - Product User Guide Version 2.0. (2017).

71. Rutgers, M. et al. Mapping earthworm communities in Europe. Appl. Soil Ecol. 97, 98–111 (2016).

72. Delgado-Baquerizo, M. et al. Ecological drivers of soil microbial diversity and soil biological networks in the Southern Hemisphere. Ecology 99, 583–596 (2018).

73. Chen, S., Zou, J., Hu, Z., Chen, H. & Lu, Y. Global annual soil respiration in relation to climate, soil properties and vegetation characteristics: Summary of available data. Agric. For. Meteorol. 198–199, 335–346 (2014).

74. Zhang, D., Hui, D., Luo, Y. & Zhou, G. Rates of litter decomposition in terrestrial ecosystems: global patterns and controlling factors. J Plant Ecol 1, 85–93 (2008).

75. Cameron, E. et al. Global mismatches in aboveground and belowground biodiversity. Conserv. Biol.

76. Menegotto, A. & Rangel, T. F. Mapping knowledge gaps in marine diversity reveals a latitudinal gradient of missing species richness. Nat. Commun. 9, 4713 (2018).

77. Eisenhauer, N. et al. Priorities for research in soil ecology. Pedobiologia 63, 1–7 (2017).

78. Hallmann, C. A. et al. More than 75 percent decline over 27 years in total flying insect biomass in protected areas. PLoS One 12, e0185809 (2017).

79. Dirzo, R. et al. Defaunation in the Anthropocene. Science 345, 401–406 (2014).

80. Cardinale, B. J. et al. Biodiversity loss and its impact on humanity. Nature 489, 326–326 (2012).

81. Titeux, N. et al. Biodiversity scenarios neglect future land-use changes. Glob. Chang. Biol. 22, 2505–2515 (2016).

82. Popp, A. et al. Land-use futures in the shared socio-economic pathways. Glob. Environ. Change 42, 331–345 (2017).

83. Dai, A. Increasing drought under global warming in observations and models. Nat. Clim. Chang. 3, 52 (2012).

84. Kharin, V. V., Zwiers, F. W., Zhang, X. & Hegerl, G. C. Changes in Temperature and Precipitation Extremes in the IPCC Ensemble of Global Coupled Model Simulations. J. Clim. 20, 1419–1444 (2007).

85. Parmesan, C. & Yohe, G. A globally coherent fingerprint of climate change impacts across natural systems. Nature 421, 37–42 (2003).

86. Bronselaer, B. et al. Change in future climate due to Antarctic meltwater. Nature 564, 53–58 (2018).

87. Collins, M., Knutti, R., Arblaster, J., Dufresne, J. L. & Fichefet, T. Long-term climate change: projections, commitments and irreversibility. (2013).

88. Delgado-Baquerizo, M., Eldridge, D. J., Hamonts, K. & Singh, B. K. Ant colonies promote the diversity of soil microbial communities. ISME J. (2019). doi:10.1038/s41396-018-0335-2

89. Wardle, D. A. et al. Ecological linkages between aboveground and belowground biota. Science 304, 1629–1633 (2004).

90. Thomson, S. A. et al. Taxonomy based on science is necessary for global conservation. PLoS Biol. 16, e2005075 (2018).

91. Drew, L. W. Are We Losing the Science of Taxonomy? Bioscience 61, 942–946 (2011).

92. Paknia, O., Sh., H. R. & Koch, A. Lack of well-maintained natural history collections and taxonomists in megadiverse developing countries hampers global biodiversity exploration. Org. Divers. Evol. 15, 619–629 (2015).

93. Prathapan, K. D. et al. When the cure kills-CBD limits biodiversity research. Science 360, 1405–1406 (2018).

94. Neumann, D. et al. Global biodiversity research tied up by juridical interpretations of access and benefit sharing. Org. Divers. Evol. 18, 1–12 (2017).

95. Leimu, R. & Koricheva, J. What determines the citation frequency of ecological papers? Trends Ecol. Evol. 20, 28–32 (2005).

96. Hugerth, L. W. & Andersson, A. F. Analysing Microbial Community Composition through Amplicon Sequencing: From Sampling to Hypothesis Testing. Front. Microbiol. 8, 1561 (2017).

97. Terrat, S. et al. Meta-barcoded evaluation of the ISO standard 11063 DNA extraction procedure to characterize soil bacterial and fungal community diversity and composition. Microb. Biotechnol. 8, 131–142 (2015).

98. Kõljalg, U., Larsson, K. H. & Abarenkov, K. UNITE: a database providing web-based methods for the molecular identification of ectomycorrhizal fungi. New (2005).

99. Mathieu, J., Caro, G. & Dupont, L. Methods for studying earthworm dispersal. Appl. Soil Ecol. 123, 339–344 (2018).

100. Pauchard, N. Access and Benefit Sharing under the Convention on Biological Diversity and Its Protocol: What Can Some Numbers Tell Us about the Effectiveness of the Regulatory Regime? Resources 6, 11 (2017).

101. Saha, S., Saha, S. & Saha, S. K. Barriers in Bangladesh. Elife 7, (2018).

102. Prathapan, K. D. & Rajan, P. D. Biodiversity access and benefit-sharing: weaving a rope of sand. Curr. Sci. 100, 290–293 (2011).

103. Frostegård, Å., Tunlid, A. & Bååth, E. Microbial biomass measured as total lipid phosphate in soils of different organic content. J. Microbiol. Methods 14, 151–163 (1991).

104. Campbell, C. D., Chapman, S. J., Cameron, C. M., Davidson, M. S. & Potts, J. M. A rapid microtiter plate method to measure carbon dioxide evolved from carbon substrate amendments so as to determine the physiological profiles of soil microbial communities by using whole soil. Appl. Environ. Microbiol. 69, 3593–3599 (2003).

105. Eppo, P. M. 7/119 (1). 2013. Nematode extraction. EPPO Bull 43, 471–495

106. ISO/FDIS. Soil quality - Sampling of soil invertebrates - Part 1: Hand-sorting and extraction of earthworms. (ISO, 2018).

107. ISO. Soil quality - Sampling of soil invertebrates - Part 4: Sampling, extraction and identification of soil-inhabiting nematodes. (ISO, 09–2011).

108. Hunter, P. A DEAL for open access: The negotiations between the German DEAL project and publishers have global implications for academic publishing beyond just Germany. EMBO Rep. 19, (2018).

109. Knapp, A. K. et al. Past, Present, and Future Roles of Long-Term Experiments in the LTER Network. Bioscience 62, 377–389 (2012).

110. van der Linde, S. et al. Environment and host as large-scale controls of ectomycorrhizal fungi. Nature 558, 243–248 (2018).

111. Terrat, S. et al. Mapping and predictive variations of soil bacterial richness across France. PLoS One 12, e0186766 (2017).

112. Phillips, H. R. P. et al. Red list of a black box. Nature Ecology & Evolution 1, 0103 (2017).

113. Davison, J. et al. Microbial island biogeography: isolation shapes the life history characteristics but not diversity of root-symbiotic fungal communities. ISME J. (2018). doi:10.1038/s41396-018-0196-8

114. Overmann, J. Significance and future role of microbial resource centers. Syst. Appl. Microbiol. 38, 258–265 (2015).

115. Overmann, J. & Scholz, A. H. Microbiological Research Under the Nagoya Protocol: Facts and Fiction. Trends Microbiol 25, 85–88 (2017).

116. Bockmann, F. A. et al Brazil’s government attacks biodiversity. Science 360, 865 (2018).

117. Scbd-Unep. Nagoya Declaration on Biodiversity in Development Cooperation. 2 (UNEP, 2010).

118. Perrings, C. et al. Ecosystem services for 2020. Science 330, 323–324 (2010).

119. Bond-Lamberty, B. & Thomson, A. A global database of soil respiration data. Biogeosci. Discuss. 7, 1321–1344 (2010).

120. Bamforth, S. S. Interpreting soil ciliate biodiversity. Plant Soil 170, 159–164 (1995).

121. Mathieu, J. EGrowth: A global database on intraspecific body growth variability in earthworm. Soil Biol. Biochem. 122, 71–80 (2018).

122. Fierer, N., Strickland, M. S., Liptzin, D., Bradford, M. A. & Cleveland, C. C. Global patterns in belowground communities. Ecol. Lett. 12, 1238–1249 (2009).

123. Nelson, M. B., Martiny, A. C. & Martiny, J. B. H. Global biogeography of microbial nitrogen-cycling traits in soil. Proc. Natl. Acad. Sci. U. S. A. 113, 8033–8040 (2016).

124. Bahram, M. et al. Structure and function of the global topsoil microbiome. Nature 560, 233–237 (2018).

125. Ramirez, K. S. et al. Detecting macroecological patterns in bacterial communities across independent studies of global soils. Nature Microbiology 3, 1–8 (2017).

126. Leff, J. W. et al. Consistent responses of soil microbial communities to elevated nutrient inputs in grasslands across the globe. Proc. Natl. Acad. Sci. U. S. A. 112, 10967–10972 (2015).

127. Gilbert, J. A., Jansson, J. K. & Knight, R. The Earth Microbiome project: successes and aspirations. BMC Biol. 12, 69 (2014).

128. Fierer, N. & Jackson, R. B. The diversity and biogeography of soil bacterial communities. Proc. Natl. Acad. Sci. U. S. A. 103, 626–631 (2006).

129. Darcy, J. L., Lynch, R. C., King, A. J., Robeson, M. S. & Schmidt, S. K. Global distribution of Polaromonas phylotypes--evidence for a highly successful dispersal capacity. PLoS One 6, e23742 (2011).

130. Hendershot, J. N., Read, Q. D., Henning, J. A., Sanders, N. J. & Classen, A. T. Consistently inconsistent drivers of microbial diversity and abundance at macroecological scales. Ecology 98, 1757–1763 (2017).

131. Locey, K. J. & Lennon, J. T. Scaling laws predict global microbial diversity. Proc. Natl. Acad. Sci. U. S. A. 113, 5970–5975 (2016).

132. Lozupone, C. A. & Knight, R. Global patterns in bacterial diversity. Proc. Natl. Acad. Sci. U. S. A. 104, 11436–11440 (2007).

133. Neal, A. L. et al. Phylogenetic distribution, biogeography and the effects of land management upon bacterial non-specific Acid phosphatase Gene diversity and abundance. Plant Soil 427, 175–189 (2018).

134. Shoemaker, W. R., Locey, K. J. & Lennon, J. T. A macroecological theory of microbial biodiversity. Nat Ecol Evol 1, 107 (2017).

135. Bates, S. T. et al. Examining the global distribution of dominant archaeal populations in soil. ISME J. 5, 908–917 (2011).

136. Davison, J. et al. Global assessment of arbuscular mycorrhizal fungus diversity reveals very low endemism. Science 349, 970–973 (2015).

137. Kivlin, S. N., Hawkes, C. V. & Treseder, K. K. Global diversity and distribution of arbuscular mycorrhizal fungi. Soil Biol. Biochem. 43, 2294–2303 (2011).

138. Pärtel, M. et al. Historical biome distribution and recent human disturbance shape the diversity of arbuscular mycorrhizal fungi. New Phytol 216, 227–238 (2017).

139. Põlme, S. et al. Biogeography of ectomycorrhizal fungi associated with alders (Alnus spp.) in relation to biotic and abiotic variables at the global scale. New Phytol. 198, 1239–1249 (2013).

140. Sharrock, R. A. et al. A global assessment using PCR techniques of mycorrhizal fungal populations colonising Tithonia diversifolia. Mycorrhiza 14, 103–109 (2004).

141. Tedersoo, L. et al. Towards global patterns in the diversity and community structure of ectomycorrhizal fungi. Mol. Ecol. 21, 4160–4170 (2012).

142. Öpik, M., Moora, M., Liira, J. & Zobel, M. Composition of root-colonizing arbuscular mycorrhizal fungal communities in different ecosystems around the globe: Arbuscular mycorrhizal fungal communities around the globe. J. Ecol. 94, 778–790 (2006).

143. Öpik, M. et al. The online database MaarjAM reveals global and ecosystemic distribution patterns in arbuscular mycorrhizal fungi (Glomeromycota). New Phytol. 188, 223–241 (2010).

144. Stürmer, S. L., Bever, J. D. & Morton, J. B. Biogeography of arbuscular mycorrhizal fungi (Glomeromycota): a phylogenetic perspective on species distribution patterns. Mycorrhiza 28, 587–603 (2018).

145. Bates, S. T. et al. Global biogeography of highly diverse protistan communities in soil. ISME J. 7, 652–659 (2013).

146. Lara, E., Roussel-Delif, L. & Fournier, B. Soil microorganisms behave like macroscopic organisms: patterns in the global distribution of soil euglyphid testate amoebae. Journal of (2016).

147. Finlay, B. J., Esteban, G. F., Clarke, K. J. & Olmo, J. L. Biodiversity of terrestrial protozoa appears homogeneous across local and global spatial scales. Protist 152, 355–366 (2001).

148. Chao, A., C. Li, P., Agatha, S. & Foissner, W. A statistical approach to estimate soil ciliate diversity and distribution based on data from five continents. Oikos (2006).

149. Foissner, W. Global soil ciliate (Protozoa, Ciliophora) diversity: a probability-based approach using large sample collections from Africa, Australia and Antarctica. Biodiversity & Conservation 6, 1627–1638 (1997).

150. Nielsen, U. N. et al. Global-scale patterns of assemblage structure of soil nematodes in relation to climate and ecosystem properties: Global-scale patterns of soil nematode assemblage structure. Glob. Ecol. Biogeogr. 23, 968–978 (2014).

151. Wu, T., Ayres, E., Bardgett, R. D., Wall, D. H. & Garey, J. R. Molecular study of worldwide distribution and diversity of soil animals. Proc. Natl. Acad. Sci. U. S. A. 108, 17720–17725 (2011).

152. Robeson, M. S. et al. Soil rotifer communities are extremely diverse globally but spatially autocorrelated locally. Proc. Natl. Acad. Sci. U. S. A. 108, 4406–4410 (2011).

153. Wall, D. H. et al. Global decomposition experiment shows soil animal impacts on decomposition are climate-dependent. Glob. Chang. Biol. 13, (2008).

154. Pachl, P. et al. The tropics as an ancient cradle of oribatid mite diversity. Acarologia 57, 309–322 (2016).

155. Dahlsjö, C. A. L. et al. First comparison of quantitative estimates of termite biomass and abundance reveals strong intercontinental differences. J. Trop. Ecol. 30, 143–152 (2014).

156. Briones, M. J. I., Ineson, P. & Heinemeyer, A. Predicting potential impacts of climate change on the geographical distribution of enchytraeids: a meta-analysis approach. Glob. Chang. Biol. (2007).

157. Silver, W. L. & Miya, R. K. Global patterns in root decomposition: comparisons of climate and litter quality effects. Oecologia 129, 407–419 (2001).

158. Zhang, T. ‘an, Chen, H. Y. H. & Ruan, H. Global negative effects of nitrogen deposition on soil microbes. ISME J. 12, 1817–1825 (2018).

159. Sinsabaugh, R. L., Turner, B. L. & Talbot, J. M. Stoichiometry of microbial carbon use efficiency in soils. Ecological (2016).

160. Xu, M. & Shang, H. Contribution of soil respiration to the global carbon equation. J. Plant Physiol 203, 16–28 (2016).

161. Raich, J. W. & Tufekciogul, A. Vegetation and soil respiration: Correlations and controls. Biogeochemistry 48, 71–90 (2000).

162. Wang, J., Chadwick, D. R., Cheng, Y. & Yan, X. Global analysis of agricultural soil denitrification in response to fertilizer nitrogen. Sci. Total Environ. 616–617, 908–917 (2018).

163. Rahmati, M. et al. Development and analysis of the Soil Water Infiltration Global database. Earth System Science Data 10, 1237–1263 (2017).

164. Serna-Chavez, H. M., Fierer, N. & van Bodegom, P. M. Global drivers and patterns of microbial abundance in soil: Global patterns of soil microbial biomass. Glob. Ecol. Biogeogr. 22, 1162–1172 (2013).

165. Howison, R. A., Olff, H., Koppel, J. & Smit, C. Biotically driven vegetation mosaics in grazing ecosystems: the battle between bioturbation and biocompaction. Ecol. Monogr. (2017).

166. Lehmann, A., Zheng, W. & Rillig, M. C. Soil biota contributions to soil aggregation. Nature Ecology & Evolution 1–9 (2017).

167. Zomer, R. J., Trabucco, A., Bossio, D. A. & Verchot, L. V. Climate change mitigation: A spatial analysis of global land suitability for clean development mechanism afforestation and reforestation. Agric. Ecosyst. Environ. 126, 67–80 (2008).

168. van Straaten Oliver, Z. R. T. A. & Bossio, D. Carbon, land and water: A global analysis of the hydrologic dimensions of climate change mitigation through afforestation / reforestation. (IWMI, 2006).

169. Chen, J., Yang, S. T., Li, H. W., Zhang, B. & Lv, J. R. Research on Geographical Environment Unit Division Based on the Method of Natural Breaks (Jenks). ISPRS - International Archives of the Photogrammetry, Remote Sensing and Spatial Information Sciences XL-4/W3, 47–50 (2013).

170. Dinerstein, E. et al. An Ecoregion-Based Approach to Protecting Half the Terrestrial Realm. Bioscience 67, 534–545 (2017).

