## Supplemental Material for "Blind spots in global soil biodiversity and ecosystem function research"

### 5. Methods

#### 5.1. Literature selection and data processing

We created a dataset by collecting published literature on macroecological studies of soil biodiversity and ecosystem functions. For this literature search, we conducted a search in the Web of Science database in November 2018 within papers published between 1945 and 2018 using the keywords: (Global* OR Continental OR latitud*) AND (soil* OR belowground) AND (*function* OR *diversity OR organism* OR biota OR animal* OR invert* OR fauna*) AND distribution AND (*mycorrhiz* OR microb* OR nematod* OR bacteria* OR ant* OR fung* OR invertebrate* OR earthworm* OR protist* OR eukaryot* OR collembola* OR rotifer* OR archaea OR formic* OR mite* OR arthropod* OR respiration OR decomposition OR nitrogen-cycling OR nutrient cycling OR water infiltration OR aggregate* OR bioturbation OR biomass). These keywords were selected to encompass the maximum number of published studies, which often use a variety of expressions for describing soil biodiversity and function. Additionally, we included studies found in the references of the papers returned by the database, as well as opportunistically gathering additional studies, such as from personal bibliographic databases of global soil studies.

The initial Web of Science search returned 1186 studies, which were screened for the following three inclusion criteria: (1) studies dealing with soil taxa and/or soil ecosystem functions; (2) studies spanning more than one continent (the Pacific islands not included in any particular continent were counted as an individual continent, contributing to the inclusion of some studies); and (3) studies that span across an entire continent (i.e., a study focussing only on Europe would not be included but if larger scales would be assessed - e.g., across Eurasia - the study would be included). From these, the number of studies was reduced to 58 that, after being complemented with additional studies, resulted in a dataset with 62 studies dealing with processes and patterns of soil biodiversity and/or soil ecosystem function across global gradients ranging from 1995[120] to 2018[121] (Figure M1, Table 2).

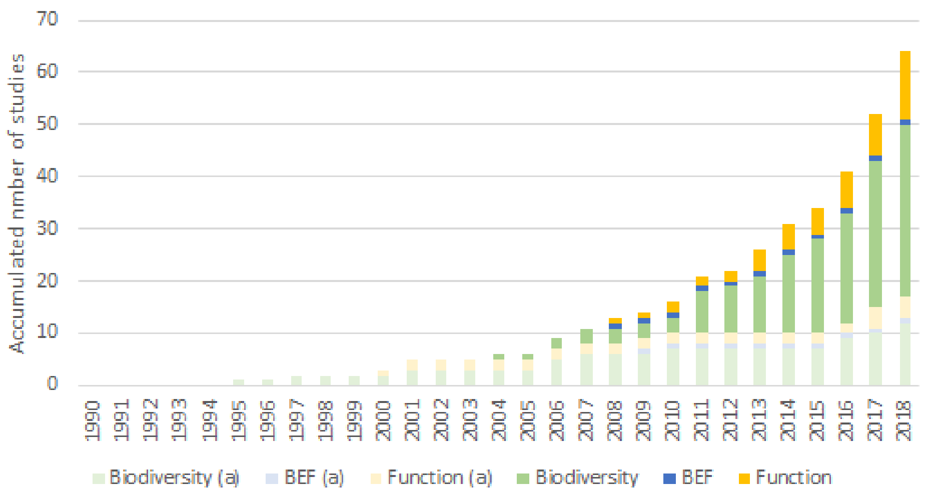

**Figure M1** Accumulated number of papers screened for this analysis (see Table 2). Studies were classified in soil biodiversity, function or biodiversity and ecosystem function (BEF), according to the subject of the study. (a) corresponds to the number of studies that were not included due to the underlying data not being suited for this analysis (e.g., based on national level information) or data availability issues. Overall, ~72.6% of the total number of studies identified were included in the analysis ranging from 2004 to 2018.

The 62 studies selected were then classified according to the taxon and/or function that was subject of the paper and screened for the availability of point coordinates for the sampling sites underlying the study (Table 2). We obtained the locations of the sampling sites that supported each manuscript, either by using published data or by contacting the corresponding authors of the publications. Papers including only national or regional level information were not included in this analysis. This was not done because of any consideration about the quality of the study nor related to bad reporting, but rather because of the lack of comparable data across studies. Overall, ~72.6% (45 studies) of the total number of studies identified were included in the analysis (see Table 2 for more details). This allowed us to cover ten different taxa and five (from a total of seven) ecosystem functions with important relevance for biogeochemical cycles[2,122,123] and the supply of key ecosystem services[1,3] (Fig. 1; Table 2).

**Table 2** List of studies included in the current assessment according to the different taxa and functions. Given the thematic scope of some studies, an individual study can be included in more than one taxon/function.

| **Soil Biodiversity** | **Included studies** | **Not included studies** |
|---|---|---|
| Bacteria | 10,14,35,63,72,123–130 | 122,131–134 |
| Archaea | 72,123,126,127,135 | 131,134 |
| Fungi | 42,113,126,127,130,136–141 | 122,131,142–144 |
| Protista | 72,145,146 | 120,147–149 |
| Nematoda | 13,150,151 | 122 |
| Rotifera | 151,152 | |
| Collembola | 153 | 122 |
| Acari | 151,153 | 122,154 |
| Formicoidea | 34,43,155 | |
| Oligochaeta | 43,121,151,153,156 | |
| **Soil Functions** | | |
| Decomposition | 64,74,153 | 157 |
| Soil Respiration | 60,73,119,158–160 | 161 |
| Nutrient Cycling | 162 | |
| Water Infiltration | 163 | |
| Secondary Productivity | 158,164 | 122 |

| | | |
|---|---|---|
| Bioturbation | | [165] |
| Soil Aggregate Stability | | [166] |

Available pairs of coordinates were then georeferenced and projected to WGS84 to create a dataset comprising all individual studies. The geographic coordinates of the sites were included based on the data provided by the studies and no spatial corrections were made (Fig. 1). As a final step, sampling sites located outside of the terrestrial scope of this paper - which excludes Antarctica and Greenland - were removed from the subsequent analysis.

### 5.2. Completeness and estimation of global representation

One of the main objectives of this paper is to describe the ability of current soil macroecological research to capture the diversity of conditions affecting the soil realm, here described as the bulk of characteristics encompassing the soil (including physical and chemical properties), climate (including properties that affect soil conditions e.g., related to soil humidity and temperature), geomorphology (including global topographic properties), and aboveground diversity[1,40]. With this definition, we identified 15 environmental and diversity descriptor variables of the soil realm (Table 3) that we used to characterize how the overall sampling locations for each study capture the global environmental and aboveground diversity scope. Initial calculations were made using the original resolution of the different datasets although, for the final spatial mapping and integration, all datasets were harmonized to an ~1km$^2$ (at the equator) resolution using a resampling algorithm without changing the original values - i.e., focussing only on pixel disaggregation with a nearest neighbour classifier.

**Table 3** Environmental variables defining the soil realm.

| Environmental variable | Dataset | Reference |
|---|---|---|
| Soil Carbon | SoilGRIDS - global soil information based on automated mapping | [65] |
| Soil pH | SoilGRIDS - global soil information based on automated mapping | [65] |
| Clay content | SoilGRIDS - global soil information based on automated mapping | [65] |
| Sand content | SoilGRIDS - global soil information based on automated mapping | [65] |
| Silt Content | SoilGRIDS - global soil information based on automated mapping | [65] |
| Soil type | SoilGRIDS - global soil information based on automated mapping | [65] |
| Mean annual temperature | CHELSA - Climatologies at high resolution for the earth's land surface areas | [67] |
| Mean annual precipitation | CHELSA - Climatologies at high resolution for the earth's land surface areas | [67] |
| Temperature seasonality | CHELSA - Climatologies at high resolution for the earth's land surface areas | [67] |
| Precipitation seasonality | CHELSA - Climatologies at high resolution for the earth's land surface areas | [67] |
| Aridity | CGIAR-CSI - Global aridity database | [167,168] |

| | | |
|---|---|---|
| Potential Evapotranspiration | CGIAR-CSI - Global potential evapotranspiration database | 167,168 |
| Elevation | GMTED2010 - Global Multi-resolution Terrain Elevation Data | 68 |
| Land cover type | ESA CCI - Global Land Cover database | 70 |
| Plant diversity | Global plant diversity | 69 |

For each study, we examined how these sample-based distributions capture the diversity of global conditions with the purpose of finding the "*blind spots*" of global soil ecosystem research corresponding to underrepresented areas of low environmental and aboveground diversity representation. To do this, we compared the global histogram of each of these environmental and aboveground diversity variables (Table 3) with the one obtained using the sampling sites for every particular study. These histograms were classified using a natural breaks (Jenks) method[169], with the exception of land cover and soil type for which the original categorical classification was maintained. This procedure allowed us to identify particular ranges of environmental and diversity conditions that are overrepresented in current literature and not assume total coverage by just looking at the spatial distribution of sampling sites. For example, having one sampling site in the tropics may wrongly give the impression that tropical regions are covered when in fact this sampling site can only cover a very small range of the total tropical spectrum. Given the small scale differences in soil communities, by contrast with other aboveground taxa, this overrepresentation of specific conditions in detriment of others is of the utmost importance as it can produce significant interpretation biases and knowledge limitations.

For each study, we overlaid the spatial representation of all variables to include the mean and median distribution and the standard deviation across environmental variables. In order to have taxonomic and functional representations (e.g., for Bacteria, fungi, decomposition, etc.), we replicated the procedure by including all the studies identified for each taxon and function. In parallel, we used global biomes[170] as a spatial stratifier to understand global biases in soil biodiversity and function data and representation. Finally, to assess the representation of soil diversity and climate conditions, it is not enough to evaluate them independently of each other. Therefore, we combined the previously classified variables (the spatial distributions of each classified variable can be found in Fig. S4a-e) in three different groups: (a) land cover (including the combination of land cover, plant diversity and elevation); (b) soils (including the combination of organic carbon, sand content and pH); and (c) climate (including the combination of mean precipitation and temperature, and their seasonality). We finally overlayed these mapped combinations with the distribution data from each study in order to understand which combinations have the highest number of redundant studies, i.e., more than one study covering the same environmental combination.

As the current analysis was conducted as a global analysis without a regional segmentation (e.g., without partitioning the analysis per continent), sampling sites in one continent (e.g., sampling sites covering high altitude areas in Europe) may influence the coverage in other continents (e.g., may identify areas in North and South America as being partially covered). Although this may lead to the overrepresentation of some areas (and consequent underrepresentation of others), our main intent is to understand how current macroecological literature is capturing the diversity of global soil conditions.

### Supplementary Information

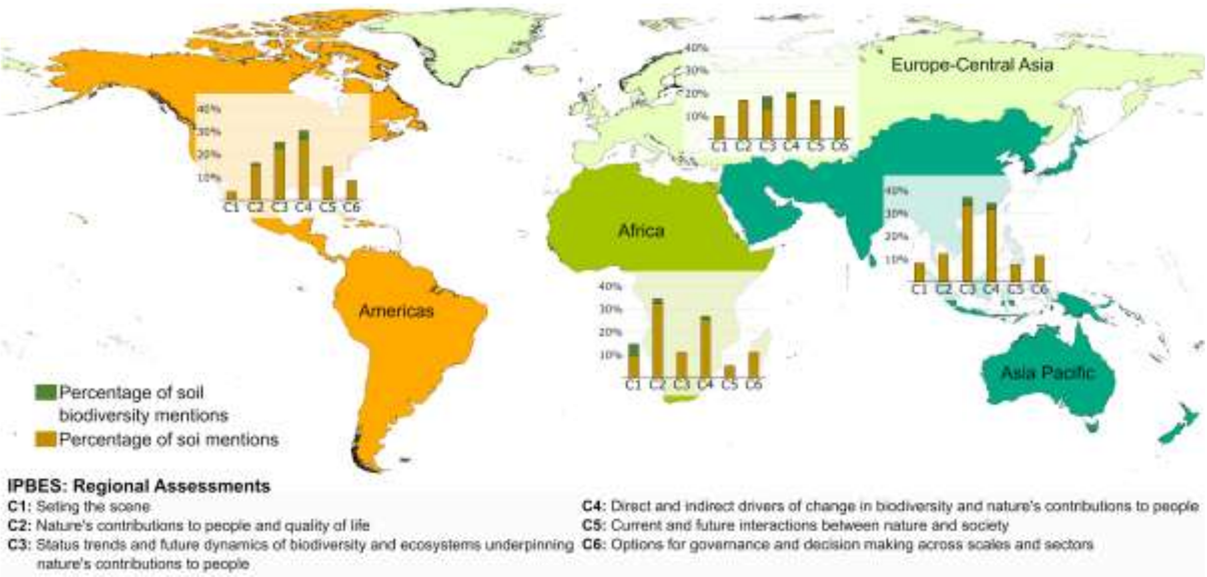


**Fig. S1** Evaluation of the extent to which the recent Intergovernmental Science-Policy Platform on Biodiversity and Ecosystem Services Regional Assessments (i.e., Americas, Africa, Europe and Central Asia, and Asia Pacific) represent soil conditions (yellow bars) and especially soil biodiversity (green bars). The stacked bar plots show the proportional representation - regarding each chapter of the assessments - of pages with references to soils, soil conditions, functions and biodiversity, in relation to the total number of pages of each chapter. The green bars refer to the proportion of pages mentioning soil biodiversity. Overall, with very few exceptions (e.g., chapter 3 of the Europe and Central Asia Assessment), most of these mentions relate to soil conditions (i.e., physical and chemical) that favour or are related to aboveground diversity, and, apart from soil carbon fixation, almost no soil functions are mentioned across the assessments. Across the assessments, soil biodiversity is systematically mentioned to highlight data gaps rather than discuss their distribution, state, or past trends. Finally, it is also clear that both biodiversity and soil ecosystem functions are almost completely absent of both chapter 5 (current and future interactions between nature and society) and from chapter 6 (options for governance and decision making across scales and sectors), two of the most fundamental chapters for decision making and the development of future conservation policies.

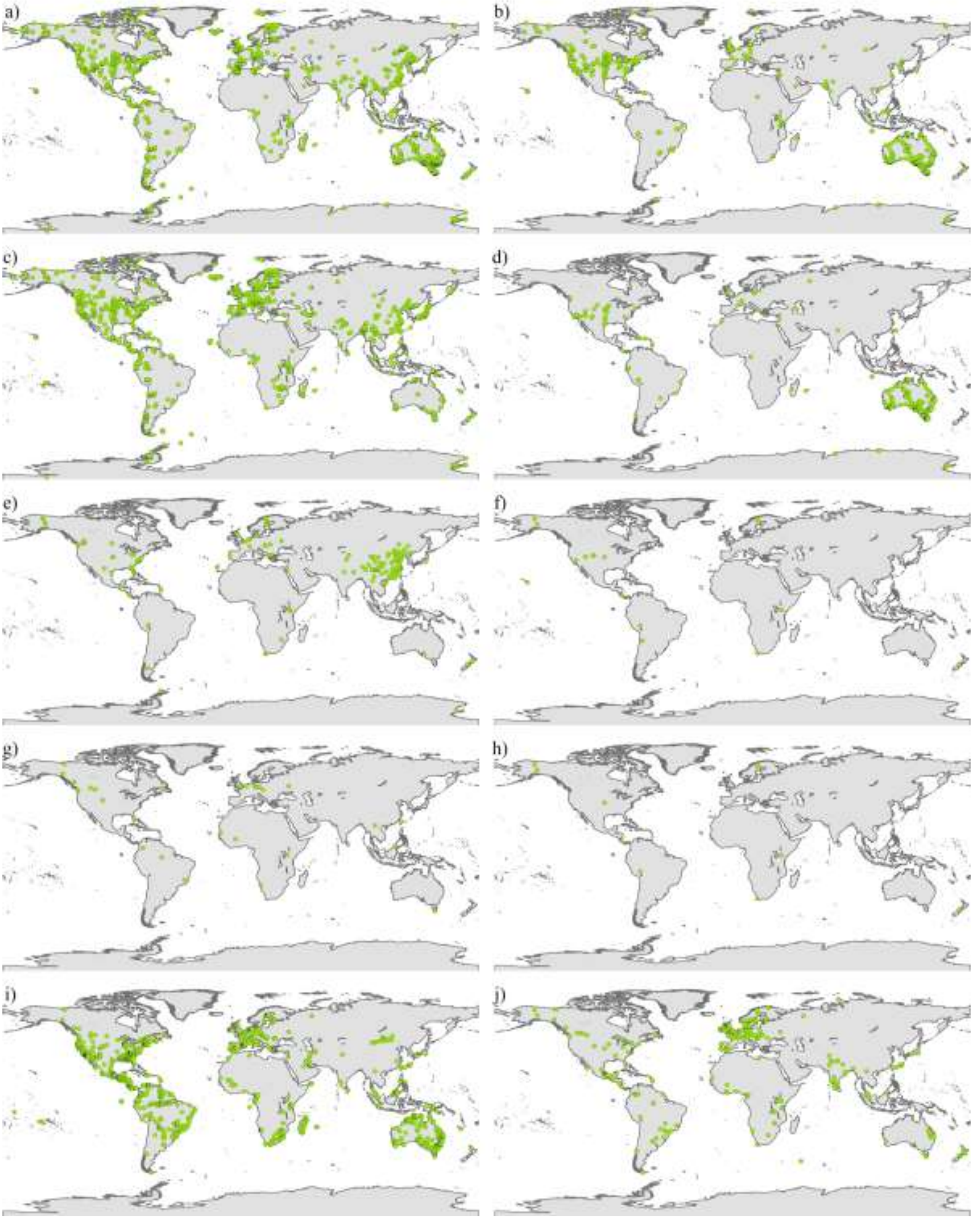


**Fig. S2a** Spatial distribution of the documented sampling sites across studies for different taxa: a) Bacteria; b) Archaea; c) fungi; d) Protista; e) Nematoda; f) Rotifera; g) Collembola; h) Acari; i) Formicoidea; and j) Oligochaeta.

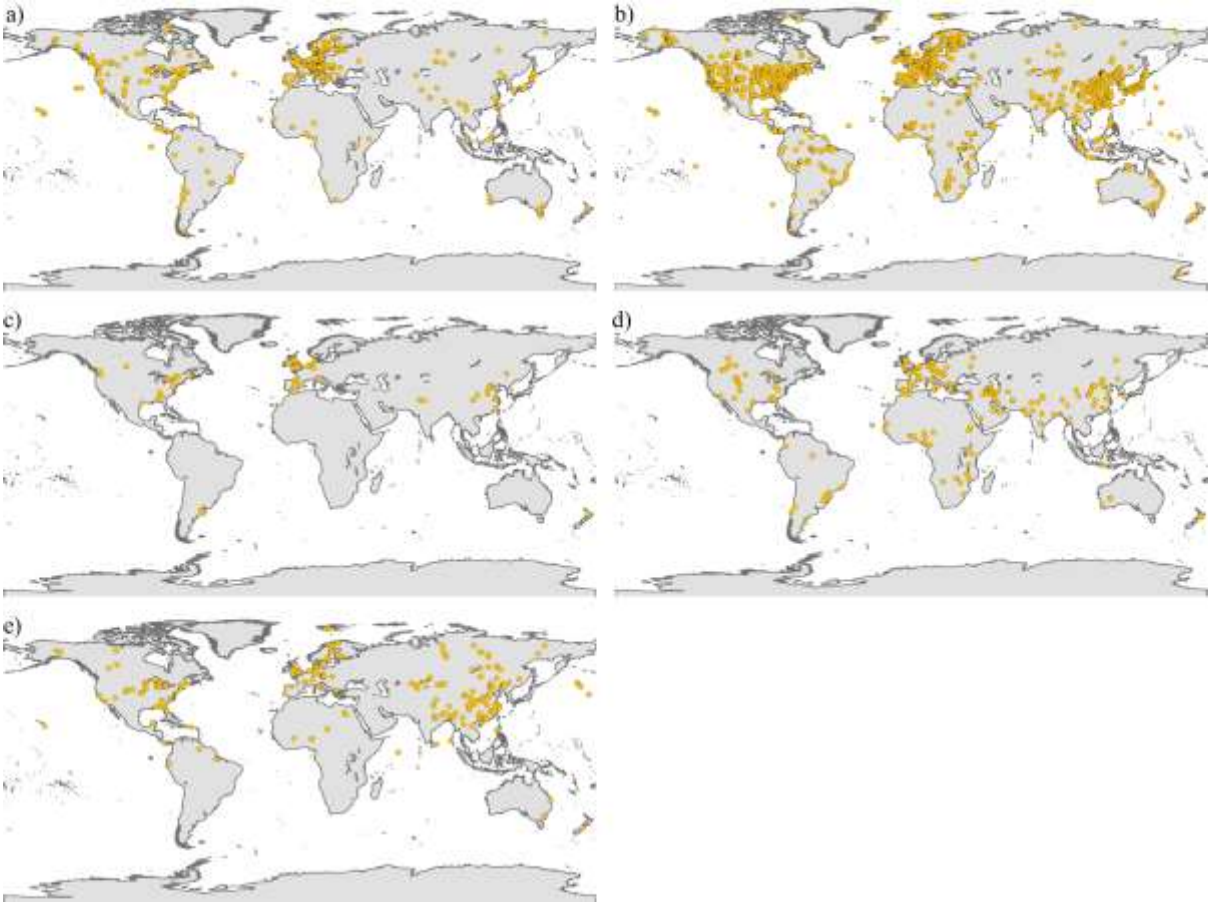


**Fig. S2b** Spatial distribution of the documented sampling sites across studies for different soil ecosystem functions: a) decomposition; b) soil respiration; c) nutrient cycling; d) water infiltration; and e) secondary productivity.

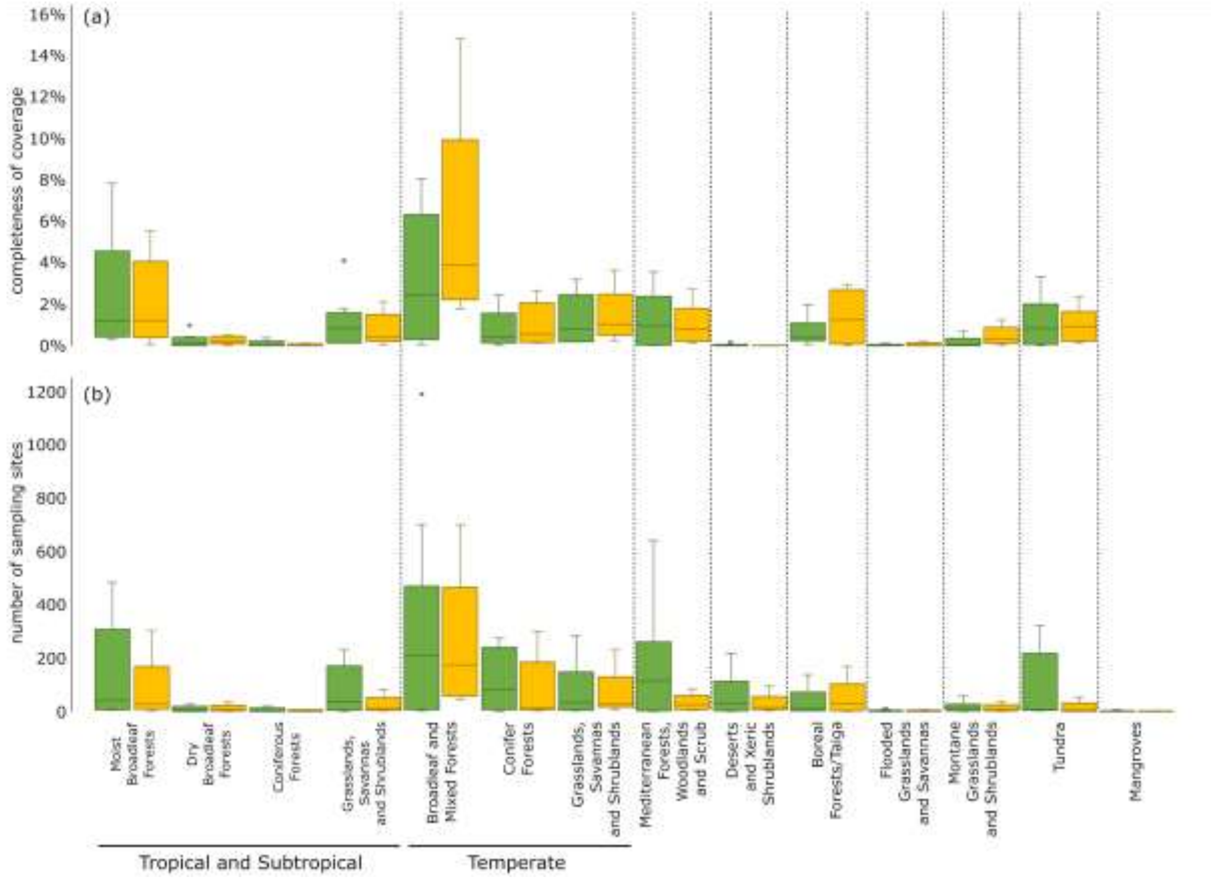


**Fig. S3** Distribution of the documented sampling sites across the different terrestrial biomes. Box plots represent the range between the different taxa (i.e., Bacteria, Archaea, fungi, Protista, Nematoda, Rotifera, Collembola, Acari, Formicoidea, and Oligochaeta; in green) and soil ecosystem functions (i.e., decomposition, soil respiration, nutrient cycling, water infiltration, and secondary productivity; in orange): (a) level of inventory spatial completeness as the percentage of 1 degree cells covered by data across biomes (the coverage extent in Mangroves was not calculated); and (b) number of sampling sites.

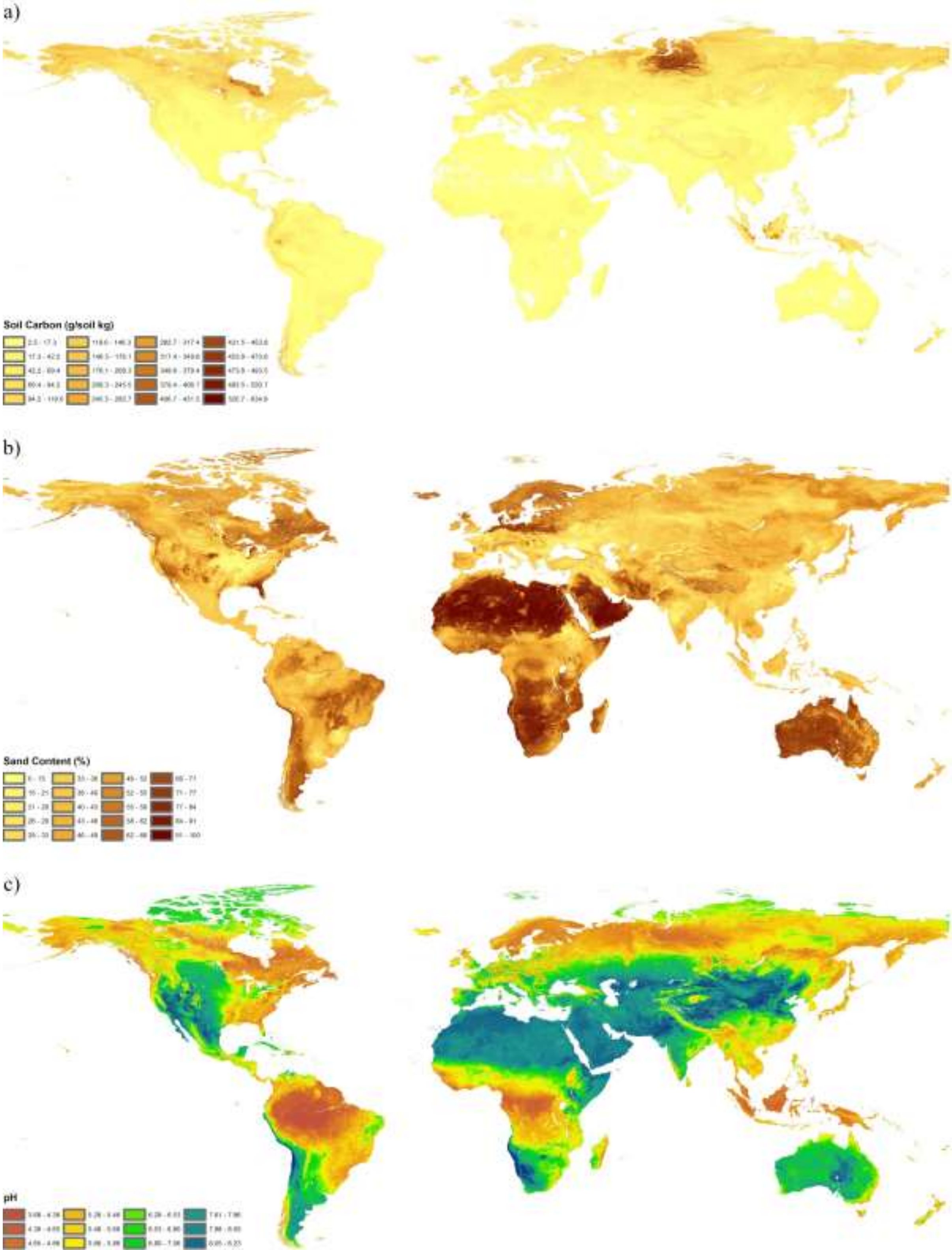


**Fig. S4a** Global distribution of the classified variables composing the soil realm: a) soil carbon[65]; b) sand content[65]; and c) pH[65].

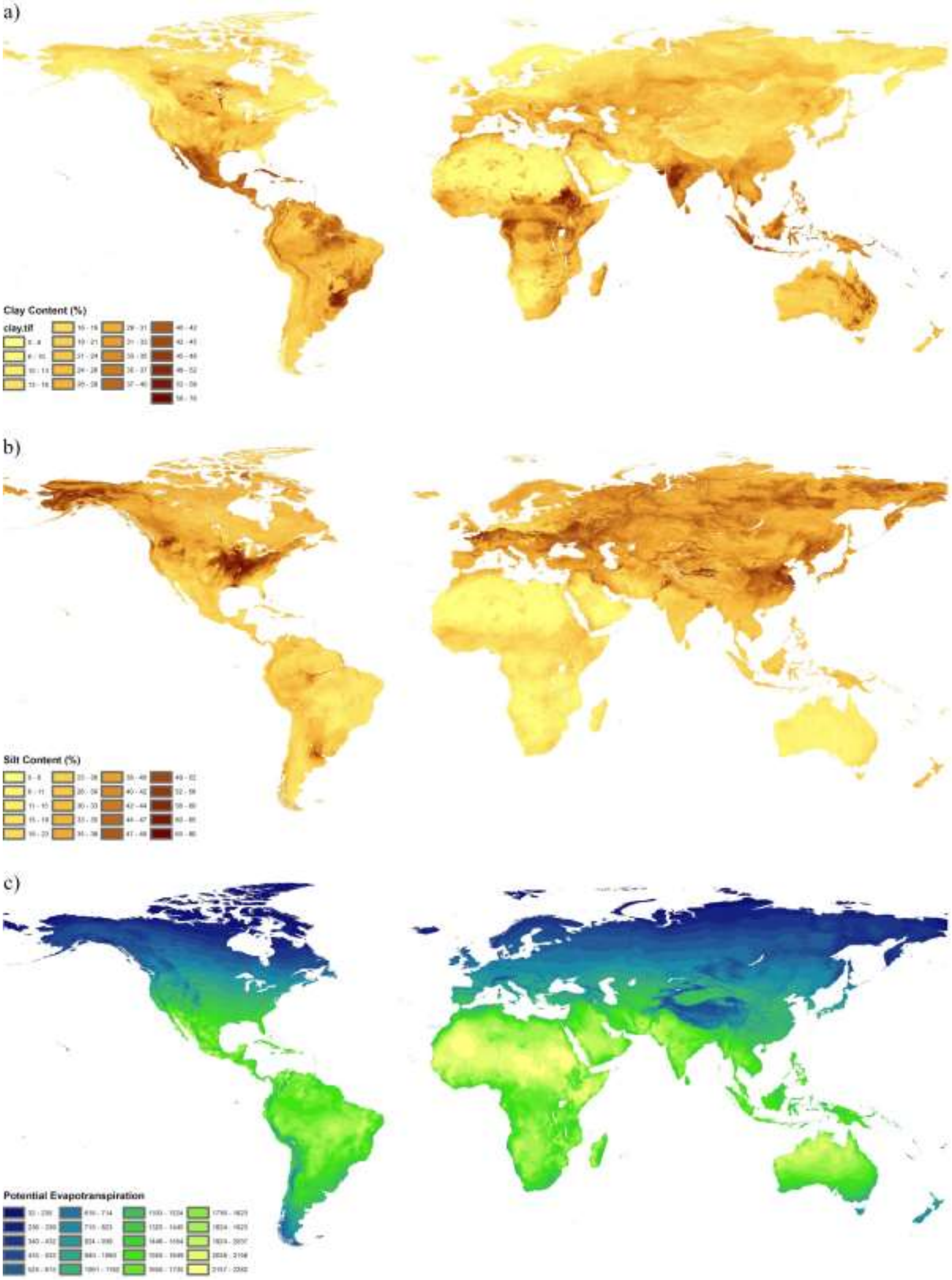


**Fig. S4b** Global distribution of the classified variables composing the soil realm: a) clay content[65]; b) silt content[65]; and c) potential evapotranspiration[167,168].

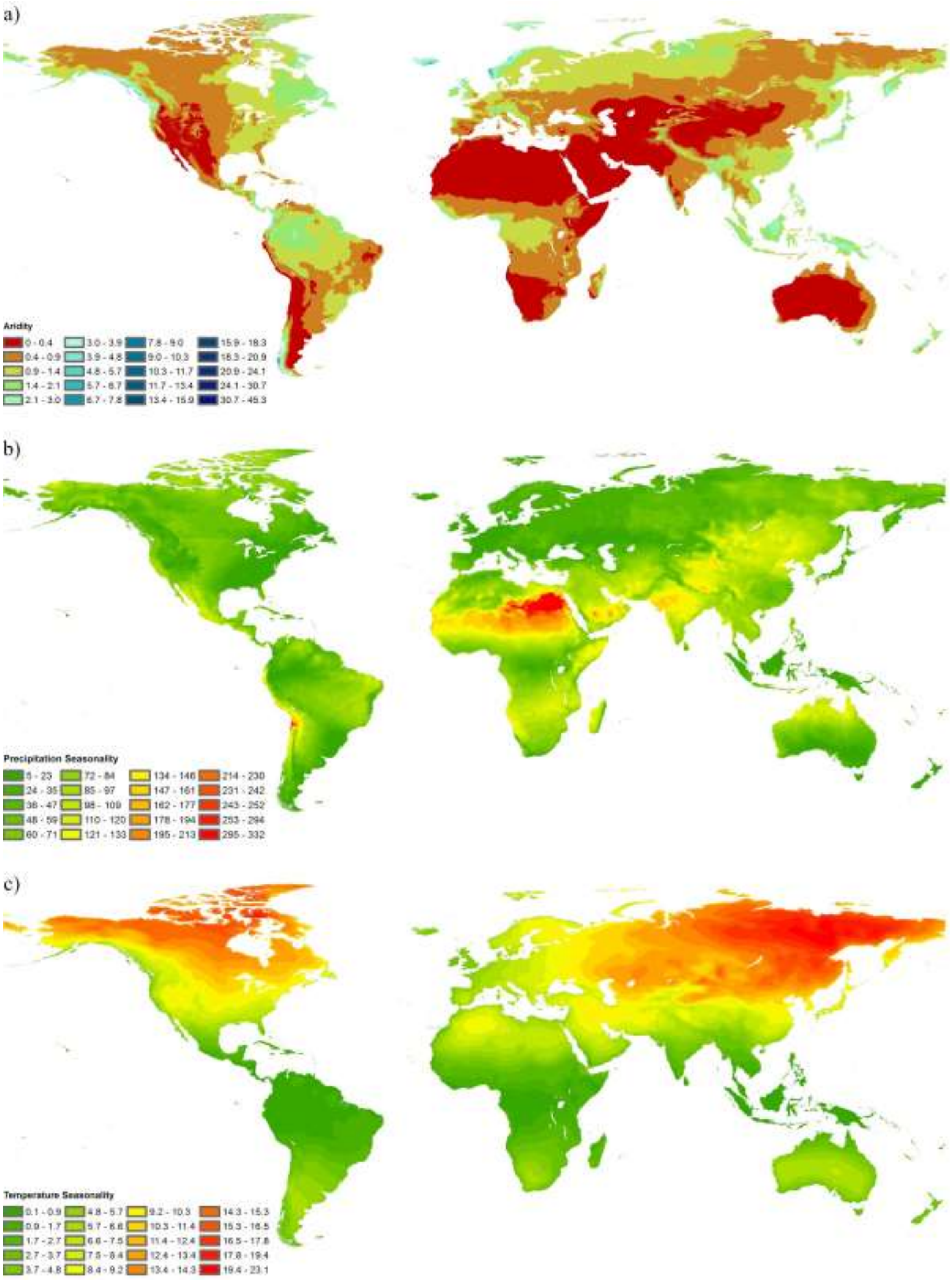


**Fig. S4c** Global distribution of the classified variables composing the soil realm: a) aridity[167,168]; b) precipitation seasonality[67]; and c) temperature seasonality[67].

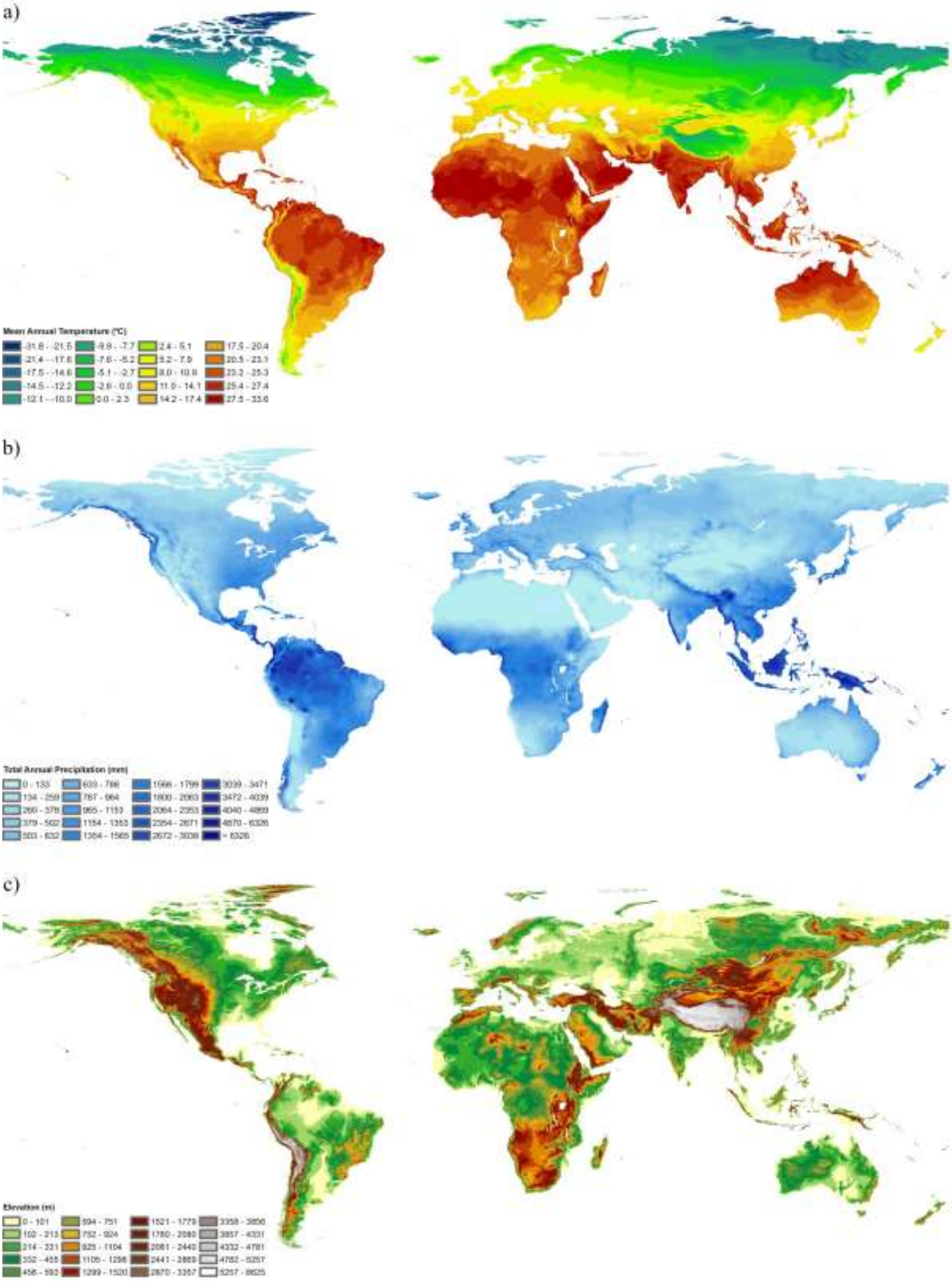


**Fig. S4d** Global distribution of the classified variables composing the soil realm: a) mean annual temperature[67]; b) total annual precipitation[67]; and c) elevation[68].

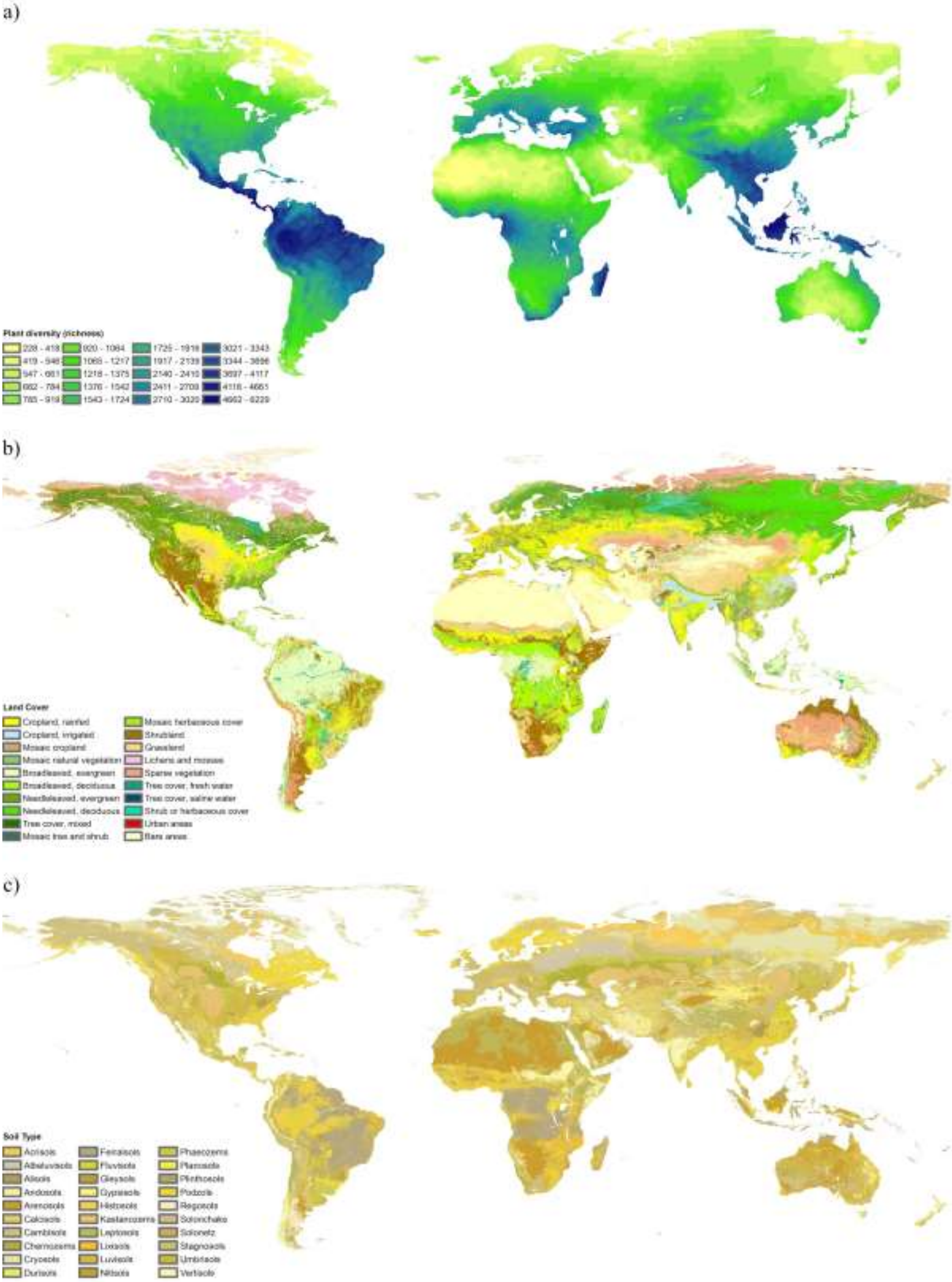


**Fig. S4e** Global distribution of the classified variables composing the soil realm: a) plant richness[69]; b) land cover[70]; and c) soil type[65].
